# A δ-cell subpopulation with a pro-β-cell identity confers efficient age-independent recovery in a zebrafish model of diabetes

**DOI:** 10.1101/2021.06.24.449706

**Authors:** Claudio A. Carril Pardo, Laura Massoz, Marie A. Dupont, David Bergemann, Jordane Bourdouxhe, Arnaud Lavergne, Estefania Tarifeño-Saldivia, Christian S. M. Helker, Didier Y. R. Stainier, Bernard Peers, Marianne L. Voz, Isabelle Manfroid

## Abstract

Restoring damaged β-cells in diabetic patients by harnessing the plasticity of other pancreatic cells raises the questions of the efficiency of the process and of the functionality of the new *Insulin*-expressing cells. To overcome the weak regenerative capacity of mammals, we used regeneration-prone zebrafish to study β-cells arising following destruction. We show that most new *in*s*ulin* cells differ from the original β-cells as they are Somatostatin+ Insulin+, but are nevertheless functional and normalize glycemia. These bihormonal cells are transcriptionally close to a subset of δ-cells in normal islets characterized by the expression of *somatostatin 1.1* (*sst1.1*), the β-cell genes *pdx1*, s*lc2a2* and *gck*, and the machinery for glucose-induced Insulin secretion. β-cell destruction triggers massive *sst1.1* δ-cell conversion to bihormonal cells. Our work shows that their pro- β-cell identity predisposes this zebrafish δ-cell subpopulation to efficient age-independent neogenesis of Insulin-producing cells and provides clues to restoring functional β-cells in mammalian diabetes models.

## Introduction

β-cells reside in pancreatic islets where they are intermingled with other endocrine cells such as α-cells producing glucagon (GCG) and δ-cells producing somatostatin (SST). Elevation of extracellular glucose concentration triggers glucose uptake by insulin-producing β-cells though the glucose transporter GLUT2 (*slc2a2*). Glucose is then metabolized to produce ATP which will cause closure of the K_ATP_ channel formed by Kir6.2 (*kcnj11)* and SUR1 *(abcc8*), membrane depolarization, Ca^2+^ influx and release through exocytosis of insulin secretory granules into the blood. In mature β-cells, this process is further amplified by other molecules such as amino acids, fatty acids, hormones (incretins GLP-1, GIP) and neural factors (dopamine, adrenaline…) via the cAMP messenger. Dysfunction of these processes leads to inadequate insulin secretion and Type 2 diabetes. Furthermore, Type 1 and type 2 diabetes are characterized by the death of β-cells as a result of autoimmune attack and failure to compensate for increased metabolic demand and chronic glucolipotoxic stress, respectively. Human adult β-cells are quiescent and barely possess the capacity to adapt to destruction through increased proliferation. Alternative mechanisms inferred from studies in mice revealed the striking plasticity of other pancreatic endocrine cell types towards the β-cell phenotype. For example, INS+ GCG+ bihormonal cells form after acute β-cell destruction mediated by transgenic expression of the diphteria toxin receptor (DTR) in adult mice (Thorel et al., 2010). These cells derive from α-cells that switched on the β-cell markers Pdx1, Nkx6.1 and Ins through direct conversion, leading to restoration of about 10% of the β-cell mass after 10 months. This process is quite slow and inefficient as only a small fraction of α- cells (1-2 %) are converted and adult DTR mice do not survive without injection of insulin during the first months after ablation. On the other hand, at juvenile stages, β-cell neogenesis occurs from massive transdifferentiation of SST producing δ-cells (Chera et al., 2014). In this case, δ-cells dedifferentiate, lose *Sst* expression, replicate and redifferentiate into β-cells. About 23% of the initial β-cell mass has recovered 4 months after ablation emphasising faster and more efficient improvement of glycemia than in adults. Hence, different mechanisms of regeneration or adaptation to loss of β-cells denote the presence of brakes inherent to adult mammals and consequent age-dependent competence.

In contrast to the limited regeneration capacity of adult mammals, adult zebrafish are notorious for their potent, spontaneous and rapid regeneration of β-cells (Delaspre et al., 2015; Ghaye et al., 2015; Moss et al., 2009). Unlike mouse models in which regeneration via progenitors or precursors in adults is debated, restoration of the lost β-cells is well recognised in adult zebrafish to involve regenerative processes from progenitor-like cells present in the ducts (Delaspre et al., 2015; Ghaye et al., 2015). In zebrafish, β-cell destruction can be accomplished using a chemo-genetic system based on the transgenic expression of the bacterial nitroreductase (NTR) under the control of the *ins* promoter where cell death is induced by a nitroaromatic prodrug (Bergemann et al., 2018; Curado et al., 2007; Pisharath, Rhee, Swanson, Leach, & Parsons, 2007). After a huge rise of glycemia within 3 days, the pancreas is replenished with new β-cells within 2 to 3 weeks which correlates to a return to normoglycemia.

*De novo* formation of β-cells in order to repair damaged islets constitutes a promising therapeutic perspective for diabetic patients. However, new β-cells could show differences in their number and identity impacting on their activity. For example, the presence in mice of GCG+ INS+ cells, though apparently functional, should be considered cautiously as inappropriate differentiation of β-cells and impaired maturation or identity are common shortcoming in diabetes ((Moin & Butler, 2019) for review).

Using the adult zebrafish as regeneration model, we investigated the identity of regenerated β-cells and discovered that most new *in*s-expressing cells are Ins+ Sst1.1+ bihormonal cells that are functional and normalize glycemia after β-cell ablation. We identified a specific δ-cell subpopulation with a transcriptomic profile presenting similarities with β-cells and which directly converts to bihormonal cells upon the loss of β-cells. Finally, we showed that p53 activity is important for Ins+ Sst1.1+ bihormonal formation.

## Results

### Most regenerated β-cells coexpress INS and SST in adult zebrafish

New β-cells are already present in the main islet within a few days after ablation in zebrafish (Delaspre et al., 2015; Ghaye et al., 2015; Moss et al., 2009). To characterize these new β-cells, we first sought to investigate whether they could express other hormones in addition to *ins* as documented for human *in vitro* models (Bruin et al., 2014; Petersen et al., 2017; Vethe et al., 2017) and murine *in vivo* models (Thorel et al., 2010). As Gcg, expressed by α-cells, have previously been reported to be unaffected after β-cell ablation in adult zebrafish (Delaspre et al., 2015), we examined the δ-cell marker Sst. β-cells were ablated in 6- to 10-month old *Tg(ins:NTR*-P2A-mCherry)* (Bergemann et al., 2018) adult fish. Basal blood glucose was monitored to evaluate ablation (3 days post treatment, dpt) and regeneration (20 dpt) and the main islet was analysed by immunofluorescence. As expected, fasted basal blood glucose measured at 3 dpt dramatically raised (510 ± 126 mg/dl) compared to CTL fish (69 ± 18 mg/dl) which reflected recent and efficient ablation (Figure 1A). After 20 days, glycemia was impressively improved though still slightly above normal values (119 ± 31 mg/dl). Immunofluorescence on control islets showed robust staining of the endogenous Ins and Sst hormones demarcating β- and δ-cells, respectively, without appreciable overlap (Figure 1B). In contrast, 20 dpt islets contained many cells with weak Ins signal that also displayed Sst immunolabelling, as well as SST-only positive cells (Figure 1B). A preliminary RNAseq experiment determined that the *somatostatin* gene expressed in *ins*+ cells after ablation was *sst1.1* (not shown). We therefore created a *Tg(sst1.1:eGFP)* reporter line driving GFP in all *sst1.1*-expressing cells which was verified to be inactive in β-cells of control islets (Figure 1-figure supplement 1). Similar to what was observed with the endogenous Sst and Ins proteins, 20 dpt islets of *Tg(sst1.1:eGFP); Tg(ins:NTR*-P2A-mCherry)* fish comprised many cells coexpressing GFP with low levels of mCherry, while control islets exhibited robust GFP and mCherry in distinct cells (Figure 1C). Strikingly, double positive cells could already be detected 3 days after ablation. In addition, while in control islets β- and δ-cells are intermingled, cells after ablation are arranged in uniform clusters composed of double positive cells.

**Figure 1.**
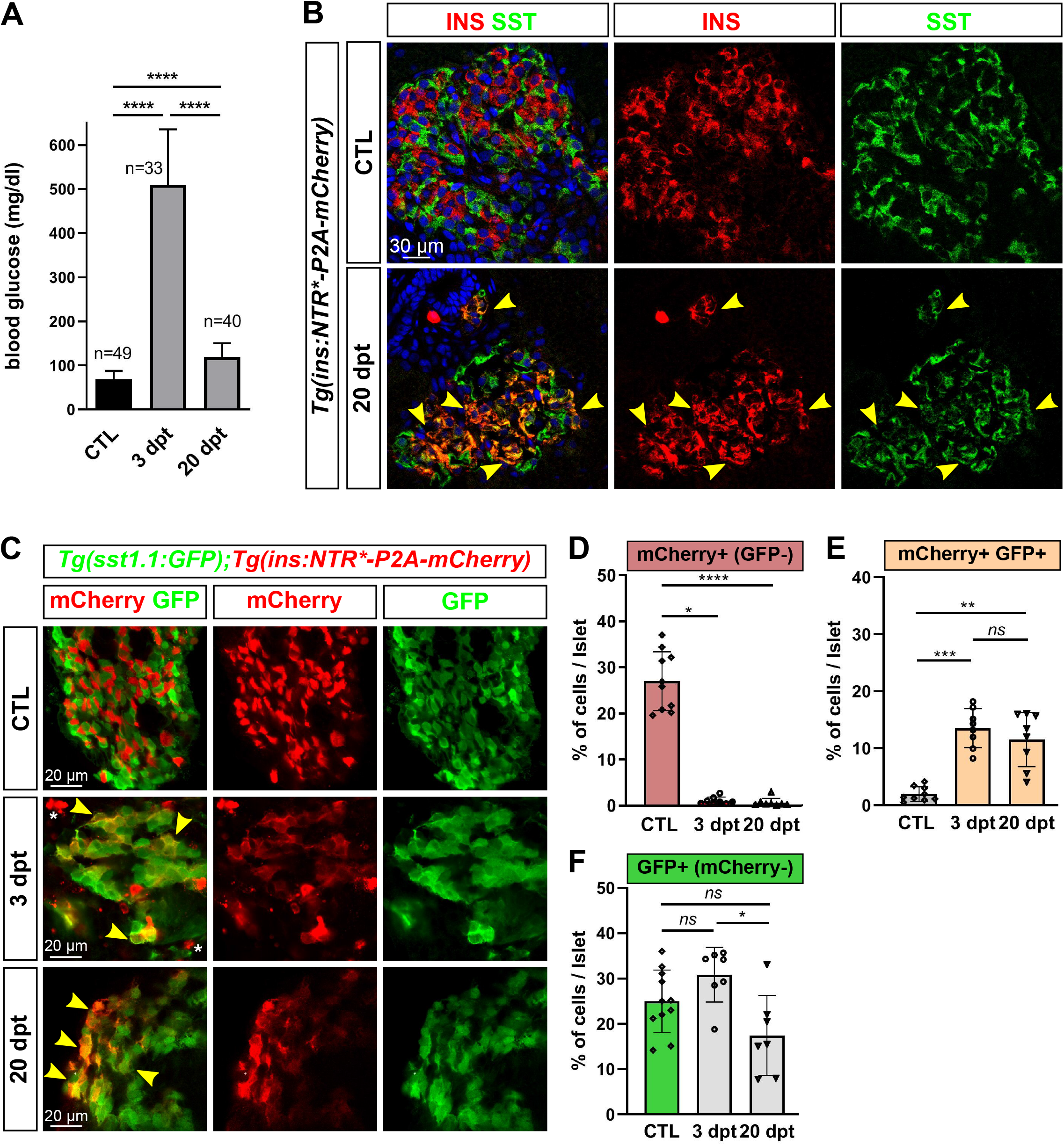
Most new *ins*+ cells after ablation and regeneration in zebrafish are INS+ SST1.1+ bihormonal cells. A) Blood glucose level (mg/ml) of adult *Tg(ins:NTR*-P2A-mCherry)^ulg034^* control fish (CTL) and 3 and 20 days post treatment (dpt) with the NFP prodrug to trigger β-cell ablation. The huge rise of glycemia at 3 dpt confirms the efficiency of ablation. One-way ANOVA Kruskal-Wallis test (with Dunn’s multiple comparisons); mean ± SD; \*\*\*\**P<0.000*1 B) Immunolabelling of β- and δ-cells with anti-INS (red) and anti-SST (green), respectively, on paraffin sections through the principal islet of *Tg(ins:NTR*-P2A-mCherry)^ulg034^* adult fish in control condition (CTL) and at 20 dpt. In CTL islet, both markers do not overlap while broad co-staining is observed at 20 dpt and represented by many yellow cells (arrowheads). C) Whole mount detection of β- and δ-cells of *Tg(sst1.1:GFP);Tg(ins:NTR*-P2A-mCherry)^ulg034^* pancreata from adult fish by immunolabelling with anti-GFP marking *sst1.1*-expressing cells and anti-mCherry for β-cells. Both cell types show no or very few overlapping in CTL fish. At 3 and 20 dpt, many double GFP+ mCherry+ cells are observed (yellow cells, arrowheads). Bright mCherry+ β-cell debris are detectable at 3 dpt (white asterisk). D-F) Proportion of GFP+, mCherry+ and GFP+ mCherry+ cells in the main islets of CTL and following β-cell ablation, based on fluorescence analysis shown in Figure 1-figure supplement 2. D) In CTL fish, mCherry+ β-cells represent 27 ± 6% of total islet cells. At 3 dpt, the fraction of mCherry+ cells per islet remaining after ablation is 1.1 ± 0.8%. E) Double mCherry+ GFP+ cells are detected at 3 and 20 dpt. These cells represent 13.5 ± 3.4% of islet cells at 3 dpt and 11.5 ± 4.8% at 20 dpt. F) GFP+ cells represent 25 ± 6.9% of control islet cells. One-way ANOVA Kruskal-Wallis test (with Dunn’s multiple comparison); *ns*, not significant, \**P<0.05*, *\*\*P<0.005, ***P<0.0005*, \*\*\*\**P<0.0001;* mean ± SD.

β-cells, *sst1.1*+ cells and double *ins*+ *sst1.1*+ cells were quantified by measuring the proportion of GFP+, mCherry+ and GFP+ mCherry+ cells in the main islet of *Tg(sst1.1:eGFP); Tg(ins:NTR*-P2A-mCherry)* adult fish (Figure 1D-F and Figure 1-figure supplement 2). In control fish, mCherry+ (GFP-) β-cells represented 27 ± 6.4% (mean ± SD) of total islet cells (Figure 1D). *sst1.1*:GFP*+* (mCherry-) cells represented 25 ± 6.9% (Figure 1F). At 3 dpt, the population of mCherry+ GFP- β-cells per islet collapsed down to 1.1 ± 0.8% of islet cells, corresponding to 4% of the initial β-cell mass. At this stage appeared a large population of double GFP+ mCherry+ fluorescent cells (13.5 ± 3.4% of islet cells) (Figure 1E). These cells account for 50% of the initial β-cell mass and 93% of the total number of *ins*-expressing cells at 3dpt. At 20 dpt, bihormonal GFP+ mCherry+ cells represented 11.5 ± 4.8% of all islet cells, still equivalent to half of the initial β-cell mass. In agreement with the weak Ins and mCherry labelling observed in bihormonal cells by immunohistochemistry, double GFP+ mCherry+ cells displayed lower mCherry fluorescence compared to β-cells of control islets (Figure 1-figure supplement 2). Importantly, mCherry+ GFP- β-cells were still a very minor population at this late time point (0.6 ± 0.9%). After ablation, GFP+ (mCherry-) cells displayed a more heterogeneous and complex profile than in CTL (Figure 1-figure supplement 2) but their abundance was not significantly altered (Figure 1F). These results indicate that *ins*+ *sst1.1*+ co-expressing cells rapidly appear in the main islet after β-cell ablation in adult fish and persist steadily for at least 20 days, and they constitute the vast majority of the new *ins*-expressing cells following ablation.

### Genesis of bihormonal cells also occurs during regeneration in larval stages and is independent of the ablation model

As in mouse the process of bihormonal cells (GCG+ INS+) formation after β-cell ablation is specific to adult stages (Thorel et al., 2010), we next asked whether bihormonal cells appear also in larvae. Indeed, we also detected bihormonal cells at 6 days post fertilization (dpf) following ablation performed at 3 dpf (Figure 2A-B) demonstrating that there is no specific competent stage for the formation of *ins+ sst1.1+* cells.

**Figure 2.**
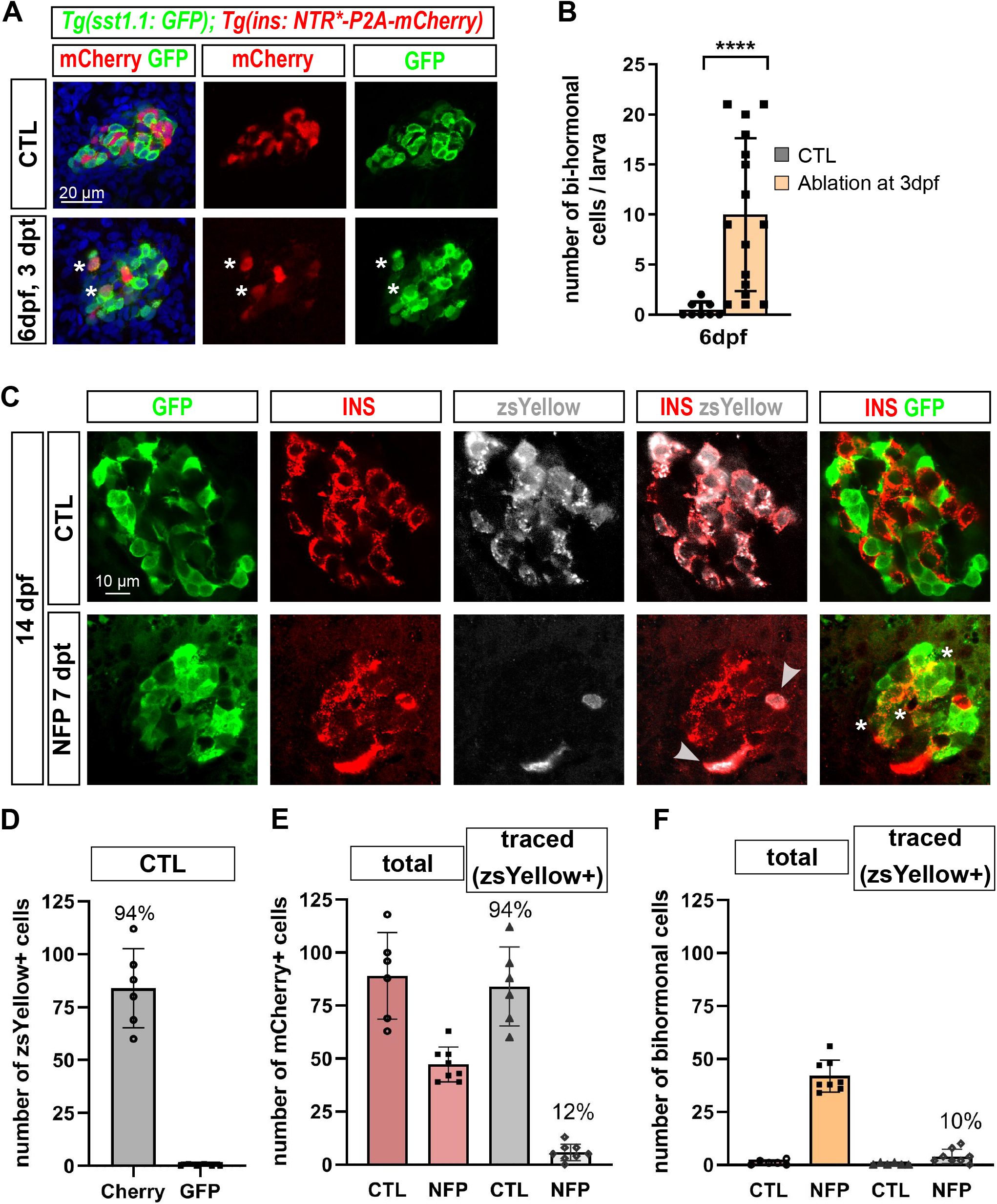
Bihormonal cell formation is age-independent and ablation model-independent. A) Representative confocal images (z-planes) showing immunodetection of β-cells (mCherry, red), *sst1.1*-expressing cells (GFP, green) and bihormonal cells (asterisks) in the principal islet 3 days (dpt) after NFP treatment at 3 dpf *Tg(sst1.1:GFP); Tg(ins:NTR*-P2A-mCherry)^ulg034^* larvae. B) Quantification of bihormonal cells co-labelled by mCherry and GFP based on confocal images. Unpaired two-tailed t-test (with Welch correction); \**P<0.05*; mean ± SD. C-F) β-cell tracing with *Tg(ins:CRE-ERT2); Tg(ubb:loxP-CFP-loxP-zsYellow); Tg(sst1.1:GFP); Tg(ins:NTR-P2A-mCherry)* larvae. CRE recombination was induced by treatment with 4-OHT treatment performed at 6 dpf to induce the expression of the lineage tracer zsYellow (grey) in Cherry+ β-cells (red) β-cell ablation was performed at 7 dpf and the lineage tracer was analysed in the islet at 14 dpf (7 days regeneration) (see Figure 2-figure supplement 2 for experimental design). C) Confocal images showing immunofluorescence of the main islet with GFP, zsYellow and INS antibodies. After ablation, traced β-cells are evidenced by zsYellow+ Cherry+ staining (grey arrowheads), bihormonal cells (Cherry+ GFP+) are indicated by white asterisks. D-F) Quantification (CTL, n=6; NFP, n=8) of the confocal images. D) ZsYellow was not detected in sst1.1:GFP+ δ-cells in CTL islets. D-E) In CTL non ablated islets, ZsYellow marked specifically and efficiently Cherry+ β-cells (84 ± 19 zsYellow+ mCherry+ cells), representing 94% of the total β-cells (89 ± 20 cells). E) After ablation (7 dpt), 47 ± 8 *ins*-expressing cells (Cherry+) were detected, of which 5.5 ± 4 (12%) were traced (Cherry+ zsYellow+) showing that most new *ins*+ cells did not derive from pre-existing β-cells. F) Most *ins*-expressing cells post-ablation are bihormonal (42 ± 7.5 cells) and only 10% of them (4 ± 3 cells) are traced by zsYellow demonstrating that they mostly derive from a non β-cell origin.

Then we questioned if *ins+ sst1.1+* cells can be induced using another system of β-cell destruction. Bihormonal cells were also observed with the NTR system using the commonly used metronidazole prodrug (not shown). To use a totally different system, we chose the Diphteria Toxin chain alpha (DTA) suicide transgene which has also been used to efficiently ablate β-cells (Ninov et al., 2013). Ablation was achieved in *Tg(ins:lox-mCherry-lox-DTA); Tg(ins:CRE-ERT2)* 7 dpf larvae by 4-OHT treatment then larvae were analysed at 16 dpf (Figure 2-figure supplement 1). Ins and Sst immunofluorescence revealed many co-expressing cells, indicating that formation of Ins+ Sst+ cells does not depend on the mode of ablation. Bihormonal cells could form by dedifferentiation - or altered identity - of β-cells. To exclude the possibility that bihormonal cells derive from pre-existing β-cells spared by the ablation, β-cells were traced using *Tg(ins:CRE-ERT2); Tg(ubb:loxP-CFP-loxP-zsYellow); Tg(sst1.1:GFP); Tg(ins:NTR-P2A-mCherry)* larvae (Figure 2C and Figure 2-figure supplement 2 for experimental design). In CTL larvae, 4-OHT induced the expression of the zsYellow lineage tracer specifically in Cherry+ β-cells and not in *sst1.1*:GFP+ cells (Figure 2C-D). 94% of the β-cells were labelled demonstrating a good efficiency (Figure 2C, E). In NFP-treated larvae, only a minority (6 ± 4, *ie* 12%) of all Ins+ cells expressed zsYellow and thus derived from β-cells (Figure 2E). In fact, most of the post-ablation Ins+ cells were bihormonal, of which most did not express the lineage tracer (42 ± 8, *ie* 90%) (Figure 2F), showing that the majority of bihormonal cells did not derive from pre-existing β-cells.

### ins+ sst1.1+ bihormonal cells share similarities with β- and δ-cells, and possess the basic machinery for glucose responsiveness

In order to characterize the *ins+ sst1.1+* cells observed after regeneration, we established their transcriptomic profile. Double GFP+ mCherry+ cells were isolated by FACS from the main islet of *Tg(sst1.1:eGFP); Tg(ins:NTR*-P2A-mCherry)* adult fish at 20 dpt and mCherry+ (GFP-). β-cells (mCherry+ GFP-) were obtained from age-matched transgenic control fish. We compared their RNAseq profiles and identified 887 DE genes with a higher expression in bihormonal cells and 705 DE genes higher in β-cells (Padj<0.05 and above 2- fold differential expression) (Figure 3A-B and Figure 3-Source Data 1). In accordance with a weak mCherry fluorescence of GFP+ mCherry+ cells as compared to native mCherry+ β-cells, expression of *ins* in bihormonal cells was 5-fold below its typical level in β-cells (Figure 3C). Also, as expected, the δ-cell hormone *sst1.1* was sharply overexpressed in GFP+ mCherry+ cells (209-fold) compared to its basal level in β-cells, and was even the top hormone just above *ins* (Figure 3C). All the other hormones known to be expressed in adult zebrafish endocrine cells, *sst1.2*, *sst2*, *gcga*, *gcgb* and *ghrl*, were expressed at much weaker levels in both *ins*+ populations (Figure 3C). Accordingly, Gcg protein was undetectable in the Cherry+ GFP+ cells by immunohistochemistry on whole islets (Figure 3-figure supplement 1). Collectively, these data confirm that bihormonal cells co-express high levels of two main hormones, *ins* and *sst1.1*, at both the mRNA and protein levels.

**Figure 3.**
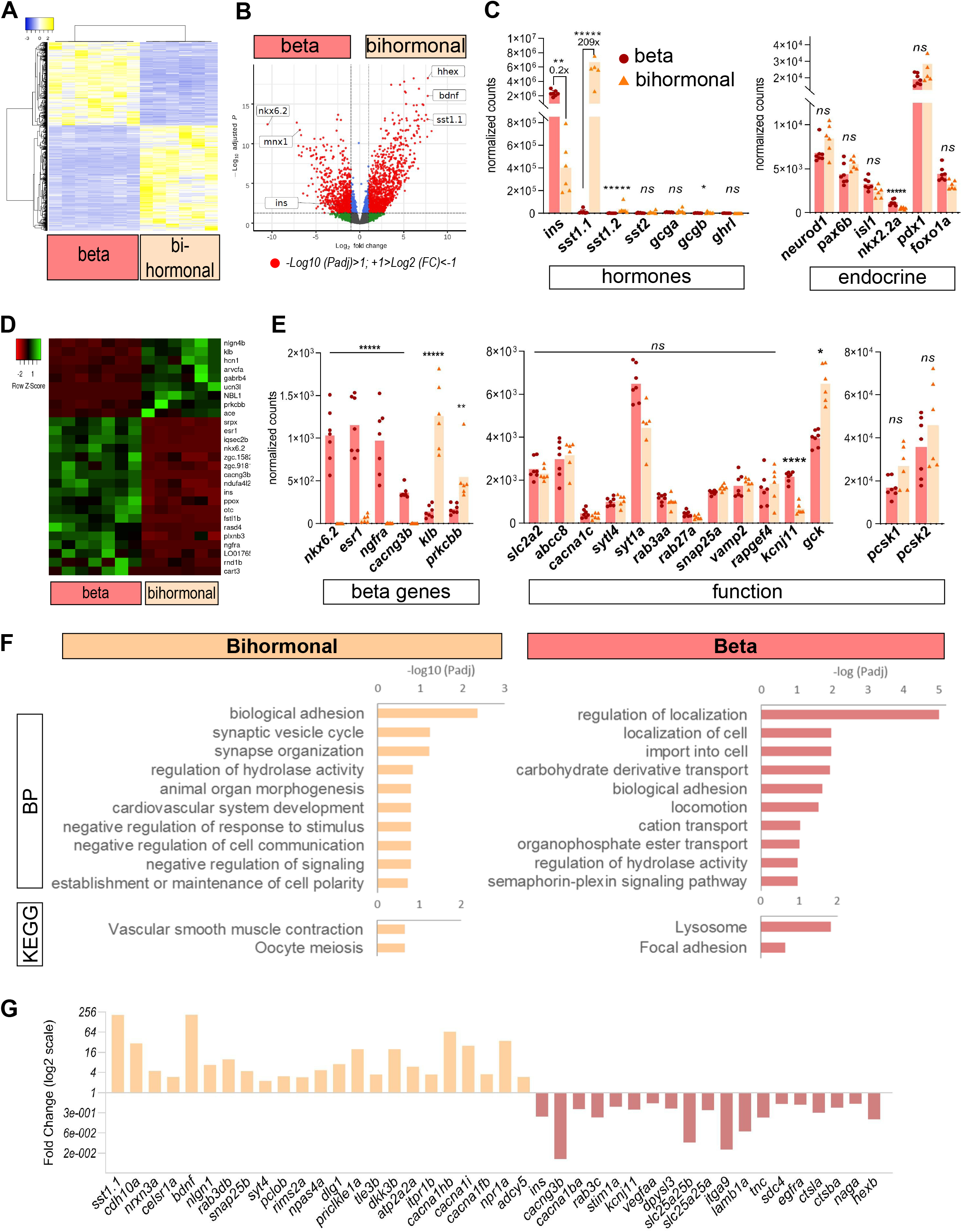
Transcriptomic comparison of bihormonal cells and β-cells. A) Heatmap representation of the transcriptomes of 20 dpt bihormonal (6 replicates) and β-cells (7 replicates) (significant DE genes). B) Volcano plot showing the distribution of genes in β-cells without ablation and bihormonal cells. The x-axis represents the log_2_ of fold change (FC) and the y-axis the log_10_ of adjusted P value (Padj) provided by DESeq. The red dots highlight the significantly DE genes (Padj<0.05). A full list of significant DE genes is provided in Figure 3-Source Data 1. C) Expression values (mean normalized reads) as provided by DESeq of the main hormones and endocrine genes in β-cell and bihormonal cell transcriptomes. *sst1.1* and *ins* are the two highest expressed hormones. Padj are calculated by DESeq. *ns:* no significant DE between the two conditions, *0.05<P*<0.005, 0.005<P**<0.0005, P*****<0.000005*. D) Heatmap plot showing the direction and amplitude of changes in expression of the β-cell markers between normal β-cells and bihormonal cells (significant DEG only). The 62 β-cell markers are provided in Figure 3-Source Data 2. E) Expression values (mean normalized reads) as provided by DESeq of selected β-cell markers and genes important for β-cell function in β-cells and bihormonal cells. Padj are calculated by DESeq. *ns:* no significant DE between the two conditions, 0.05<*<0.005, 0.005<**<0.0005, 0.00005<****<0.000005, *****<0.000005. F) Enriched Gene Ontology (GO) terms. Top 10 or Padj (FDR) <0.25 Biological Processes (BP) and KEGG pathways are shown. The plots represent the enrichment ratio of Biological Processes and KEGG pathways identified with WebGestalt (Liao et al., 2019) using the genes over- and under-expressed in bihormonal cells compared to β-cells obtained with a 2-fold differential expression and Padj<0.05. All overrepresented Biological Processes and Pathways (<FDR 0.25) are presented in Figure 3-Source Data 3 (bihormonal cells) and Figure 3-Source Data 4 (β-cells). G) Over- and under-expression of selected significantly DE genes from the BP and KEGG pathways identified in β-cells and bihormonal cells (Fold Change, log2 scale).

To further characterize these bihormonal cells, we assessed the expression of pan-endocrine genes and found that *neurod1*, *pax6b* and *isl1* showed similar expression while *nkx2.2a* was lower (Figure 3C). Importantly, *pdx1*, a β-cell determinant essential for *ins* expression, was equally expressed in both native β-cells and post-regeneration GFP+ mCherry+ cells, as well as *foxo1a*, which in mammals is a guardian of β-cell identity under stress and enhances β-cell function (Talchai, Xuan, Lin, Sussel, & Accili, 2012; Zhang et al., 2016). We thus evaluated the β-cell identity of bihormonal cells and interrogated the expression of zebrafish β-cell markers that we defined as genes enriched in β-cells (>4-fold) versus the other main pancreatic cell types, α-, *sst2* δ-cells, acinar and ductal cells, based on RNAseq data previously obtained by our group (Tarifeño-Saldivia et al., 2017) (Figure 3-Source Data 2). Among these 62 “β-cell genes” (see Methods), most (35/62) were expressed at similar levels in both *bona fide* β-cells and post-regeneration bihormonal cells. Among the 27 β-cell genes exhibiting a significant differential expression, 18 showed underexpression in bihormonal cells, of which *nkx6.2, esr1, cacng3b* and *ngfra* were almost absent (Figure 3D-E). Bihormonal cells also lacked *mnx1*, a gene important for β-cell identity in mouse (Pan, Brissova, Powers, Pfaff, & Wright, 2015). On the opposite, 9 β-cell markers showed reinforced expression in bihormonal cells such as *klb* and *prkcbb.* Of note, the pancreatic progenitor markers *nkx6.1, sox9b* and *ascl1b*, a functional orthologue of mammalian *Neurog3* (Flasse et al., 2013), were barely expressed in bihormonal cells, like in control β-cells (Figure 3-Source Data 1).

When considering key genes for β-cell function and maturation, *i.e.* glucose sensing, uptake, INS maturation and secretion, many were expressed at comparable levels in both cell types (*slc2a2, pcsk1*, *pcsk2*, *abcc8, cacna1c, chga*, *syt7a*, s*ytl4, rab3aa/ab, rab27a, snap25a, vamp2*, *rapgef4)* (Figure 3E). In contrast, the expression of *kcnj11* (encoding for the K_ATP_ channel Kir6.2) was lower while *gck* (1.6-fold), the glucose sensor necessary for coupling glucose to insulin release, was higher. *ucn3l*, a marker of mature β-cells in mammals (Blum et al., 2012) and zebrafish (Singh et al., 2017), was overexpressed in bihormonal cells.

Gene Ontology (GO) analysis of the genes overexpressed in bihormonal cells compared to β-cells showed top significant biological processes related to adhesion (*cdh10a*, *nrxn3a, celsr1a…*) and neuronal synapses (*bdnf, nlgn1, rab3db, snap25b, syt4, pclob, rims2a, npas4a*…), of which many genes are known in β-cells to be also important for Ins exocytosis. Other processes included “negative regulation of communication” (*tle3b, dkk3b*…) and “establishment or maintenance of cell polarity” (*dlg1, llgl1, prickle1a…)* (Figure 3F-G and Figure 3-Source Data 3)*. Tle* genes encode for transcriptional corepressors important for β-cell identity (Metzger, Liu, Ziaie, Naji, & Zaret, 2014). The overrepresented pathways “vascular smooth muscle contraction” *(cacna1fb, adcy5, npr1b…*) and “oocyte meiosis” (*atp2a2a, itpr1b, cacna1hb, cacna1i…)* were consistent with intracellular Calcium and cAMP signalling. Of note, *adcy5* is crucial for coupling glucose to insulin secretion in human β-cells (Hodson et al., 2014) and Npas4 is a Ca2+-dependent cytoprotective factor that blunts incretin-stimulated cAMP production and the potentiation of insulin secretion while improving glucose-mediated secretion (Sabatini et al., 2013). These data strongly suggest that bihormonal cells, like β-cells, are excitable cells that actively secrete Ins in response to glucose.

On the other side, top overrepresented processes and pathways in β-cells (Figure 3F-G and Figure 3-Source Data 4) highlighted “regulation of localization” notably with several ion channels, vesicle and cytoskeleton components (*cacng3b, cacna1ba, stim1, kcnj11, rab3c, dpysl3…), “*carbohydrate derivative transport” (*slc2a25a/b…)*, “focal adhesion” (*itga9, lamb1a, tnc, sdc4, egfra…*) and “lysosome” signature (*ctsla, ctsba, naga, hexb*…) illustrative of adult β-cells.

Altogether, these data indicate that 20 dpt bihormonal cells possess the molecular bases of functional mature β-cells such as a glucose-responsiveness and hormone secretion machinery. In line with this conclusion, these analyses did not highlight signatures of energy metabolism, implying no major differences in glucose and mitochondrial metabolism which are essential for the glucose/insulin secretion coupling. Still, bihormonal and β-cells display differential *kcnj11* expression and Ca2+ and cAMP responses demonstrating that the status and/or regulation of their secretory activity are not identical. Moreover, the respective adhesion and polarity signatures indicate differences in the way β-cells and bihormonal cells organize and communicate with their neighbours, which is evidenced by their distinct structural arrangement in the islet (see Figure 1B-C). Finally, although most β-cell genes are similarly expressed, bihormonal cells display a divergent identity such as lack of expression of the zebrafish β-cell marker *nkx6.2* and strong expression of *sst1.1* and of the δ-cell transcription factor *hhex*.

### sst1.1 δ-cells are distinct from sst2 δ-cells and display similarities with β-cells

Given the expression of *sst1.1* and *hhex* in bihormonal cells, we next sought to characterize the *sst1.1-*expressing cells in control islets. Previous transcriptomic studies of pancreatic cells by our laboratory and others detected three *Sst* genes in normal adult pancreatic islets, *sst1.1*, *sst1.2* and *sst2* (Spanjaard et al., 2018; Tarifeño-Saldivia et al., 2017). However, so far, only *sst2*-expressing cells, which also strongly express *sst1.2*, have been fully characterized (Tarifeño-Saldivia et al., 2017). We thus isolated the *sst1.1*-expressing cells from control islets of *Tg(sst1.1:eGFP)* adult fish (without ablation) to determine their transcriptomic profile. Close examination of these *sst1.1:*GFP+ cells by flow cytometry actually distinguished two subpopulations recognised by different levels of GFP fluorescence, GFP^low^ and GFP^high^ (Figure 4-figure supplement 1A), that were corroborated *in situ* by immunofluorescence on fixed whole pancreas (Figure 4A). The GFP^high^ population represented 41% of all GFP cells, *ie* 11.03 ± 2.9 % of all islet cells.

**Figure 4.**
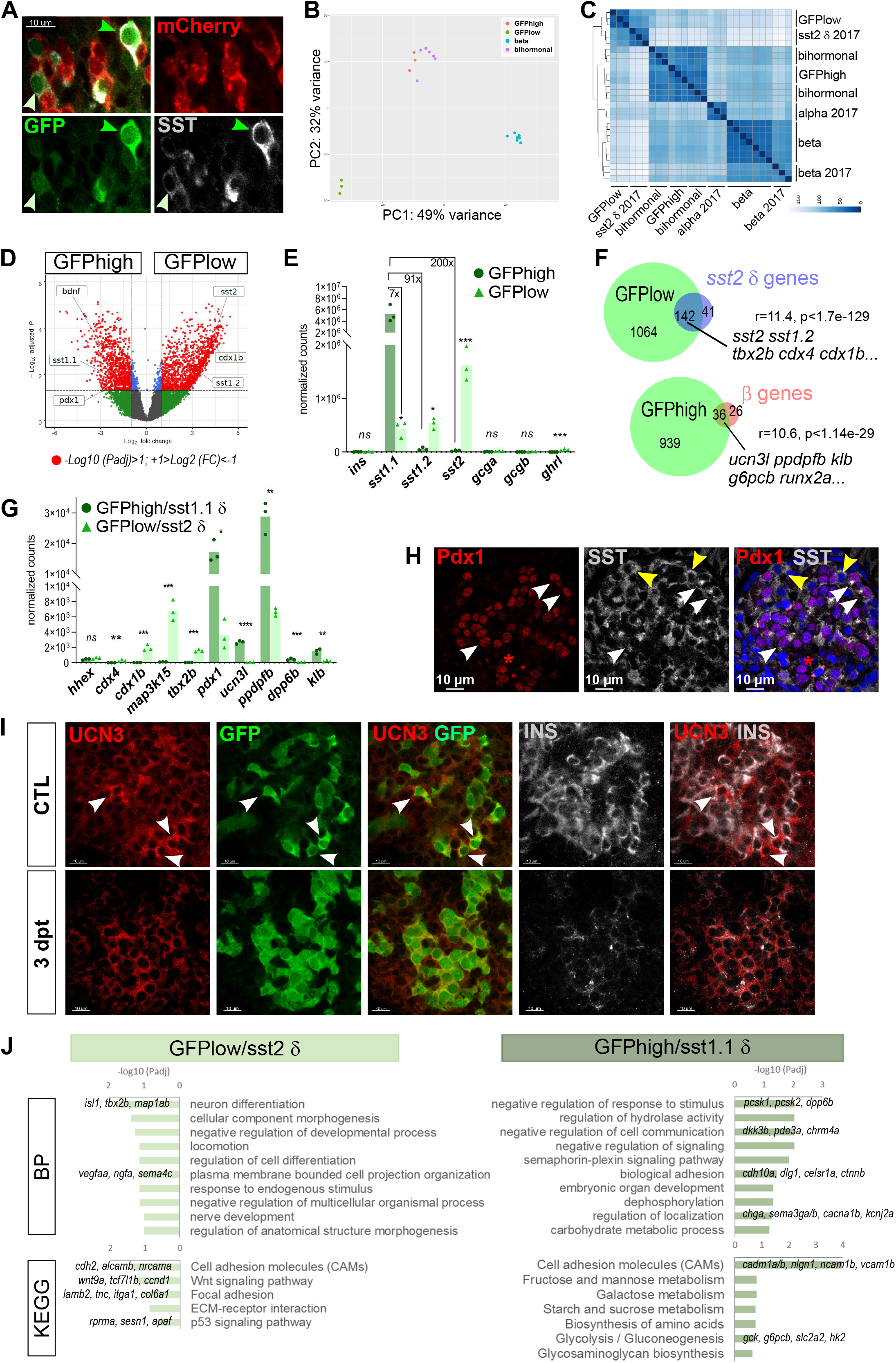
*sst1.1* δ-cells (GFP^high^) constitute a δ-cell subpopulation distinct from *sst2* δ-cells (GFP^low^) that presents similarities with β-cells. A) Immunodetection on whole mount *Tg(sst1.1:eGFP); Tg(ins:NTR*-P2A-mCherry)* islet of GFP (green), mCherry (red) and SST (gray) reveals two levels of GFP expression (green light and dark arrowheads) that parallel the expression level of SST. These cells are mCherry negative. B) PCA plot showing the separation between *sst1.1*:GFP^high^ (n=3), sst1.1:GFP^low^ (n=3), bihormonal (n=6) and β-cells (n=7) based on their transcriptomic profile. 49% of the variance is explained in PC1. PCA analysis failed to separate bihormonal and *sst1.1*:GFP^high^ cells while separated well β-cells from the sst1.1:GFP^low^ cells. The *sst1.1*:GFP^high^/bihormonal cluster located between β-cells and sst1.1:GFP^low^ cells shows that β-cells are more similar to *sst1.1*:GFP^high^/bihormonal cells. C) Heatmap plot showing the clustering of the sst1.1:GFP^high^ and sst1.1:GFP^low^ populations, the bihormonal cells, the β-cells of the present study and the previously published data for β-, α- and *sst2* δ-cells (n=3) (Tarifeño-Saldivia et al., 2017). In addition to revealing the expected clustering between both RNAseq data from β-cells ((Tarifeño-Saldivia et al., 2017) and present study), this plot also shows the clustering of the GFP^low^ cells together with *sst2* δ-cells. D) Volcano plot showing the distribution of genes expressed in GFP^high^ and GFP^low^ populations. The x-axis represents the log_2_ of fold change (FC) and the y-axis the log_10_ of adjusted P value (Padj) provided by DESeq. The list of all DE genes is provided in Figure 4-Source Data 1. E) Expression of the main pancreatic hormones in GFP^high^ and GFP^low^ populations (mean normalized reads). Expression is expressed as normalized counts and Padj are calculated by DESeq. *ns:* no significant DE between the two conditions, 0.05<*0.005, 0.005<**<0.0005, 0.0005<***<0.00005. F) Venn diagram showing the overlap between genes overexpressed in GFP^low^ cells (versus GFP^high^) and *sst2* δ-cell markers previously identified, and between genes overexpressed in GFP^high^ cells (versus GFP^low^ cells) and β-cell genes. Representation factor and P value calculated by Fisher’s exact test. The list of genes used to generate this Venn diagram is found in Figure 4-Source Data 2. G) Expression of selected β- and *sst2* δ-cell genes in each replicate of GFP^high^ and GFP^low^ cells. GFP^high^ cells distinctly express high levels of *sst1.1* and will be referred to as GFP^high^*/sst1.1* δ-cells, and GFP^low^ to GFP^low^/*sst2* δ-cells. H) Confocal images showing immunodetection of Pdx1 (anti-Pdx1, red) and SST (anti-SST, grey) on paraffin section through an adult control islet showing double Pdx1+ SST+ cells (white arrowheads) and single Pdx1- SST+ cells (yellow arrowheads). The red asterisk (*) highlight Pdx1 single positive cells β-cells. Note the particular pattern of the cellular distribution of the SST staining in Pdx1+ SST+ cells suggesting a secretory behaviour. I) Confocal images showing immunodetection of Ucn3 (red), GFP (green) and INS (grey) in CTL and 3 dpt islets from *Tg(sst1.1:eGFP); Tg(ins:NTR*-P2A-mCherry)* adult fish. In CTL islets, strong Ucn3 labelling is detected in β-cells as well as in some *sst1.1*:GFP cells (white arrowheads). After β-cell ablation, Ucn3 is principally expressed in GFP+ cells that also harbour faint INS staining. J) Biological Processes (BP) and KEGG pathways overrepresented in *sst1.1* δ-cells (UP) compared to GFP^low^ cells (DOWN) (Padj<0.25). Gene Ontology (GO) terms were identified by WebGestalt (Liao et al., 2019) using the list of DE genes between GFP^high^/*sst1.1* δ-cells and GFP^low^/*sst2* δ-cells obtained with at least 2-fold differential expression and Padj<0.05 provided by DESeq. The list of all BP and KEGG pathways below FDR 0.25 is given in Figure 4-Source Data 4 and 5.

The transcriptomic profile of these two GFP populations was obtained (Figure 4-figure supplement 1B). Principal Component Analysis (PCA) unveiled that GFP^high^ cells are more similar to β-cells than are GFP^low^ cells, and that they are also very similar to bihormonal cells (Figure 4B). In addition, clustering analysis of these two GFP populations and the other endocrine cells (β-cells, bihormonal cells, α-cells and *sst2* δ-cells from *Tg(sst2:GFP)* (Tarifeño-Saldivia et al., 2017) also showed that the GFP^high^ cells cluster together with bihormonal cells and apart from the GFP^low^ cells. Indeed, GFP^low^ cells were closer to *sst2* δ-cells than to the other endocrine subtypes (*sst1.1* δ-, α- and β-cells) (Figure 4C). Comparison of the two GFP populations identified 975 and 1206 DE genes overexpressed in GFP^high^ and GFP^low^, respectively (FC>2, Padj<0.05) (Figure 4D and Figure 4-Source Data 1). *sst1.1* was by far the predominant *sst* gene expressed in GFP^high^ cells (Figure 4E). On the opposite, *sst2* was predominant in GFP^low^ cells though these cells also expressed *sst1.2* and *sst1.1* at lower levels. In addition, while both populations expressed the universal δ-cell marker *hhex*, other previously identified markers of zebrafish *sst2* δ-cells such as *cdx4, tbx2b* and *map3k15* (Tarifeño-Saldivia et al., 2017) were specific to GFP^low^ cells (Figure 4F-G). Indeed most (3/4) of the *sst2* δ-cell genes (enriched >4-fold based on previous data (Tarifeño-Saldivia et al., 2017)) were also enriched in GFP^low^ cells (Figure F and Figure 4-Source Data 2). Ectopic activity of the *sst1.1:GFP* transgene in the *sst2* δ-cells was confirmed by ISH showing *sst2* probe signal exclusively in the weakest GFP+ cells (Figure 4-figure supplement 1C). As expected, both populations expressed only basal levels of *ins* or of the other principal hormones. These data show that the GFP^low^ cells contain *sst2* δ-cells, while the GFP^high^ population consists of a pure and distinct δ-cell population characterized by strong *sst1.1* expression.

Given the global similitude between the transcriptomes of GFP^high^*/sst1.1* δ-cells, bihormonal and β-cells, we focused on the GFP^high^*/sst1.1* δ-cells and noticed high expression of the β-cell marker *pdx1* (Figure 4G). Pdx1 immunolabelling was indeed confirmed in a subset of Sst+ cells (Figure 4H). The strong sequence conservation of the Sst1.1 peptide with human SST (Devos et al., 2002) strongly suggests that commercial anti-human SST antibodies at least recognize Sst1.1, and hence the Sst+ Pdx1+ coexpressing cells correspond to GFP^high^*/sst1.1* δ-cells. To investigate other potential similitudes of these cells with β-cells, we investigated the expression of the 62 zebrafish “β-cell genes”. Strikingly, most of the β-cell genes (36/62), such as *ucn3l, g6pcb, ppdpfb, dpp6b*…, were found enriched in GFP^high^*/sst1.1* δ-cells (Figure 4F-G and Figure 4-Source Data 2) while none was preferentially expressed in the GFP^low^ cells. By immunofluorescence, Ucn3 decorated β-cells in control islets, however, a more intense staining was detected in a subset of GFP cells presenting also robust GFP fluorescence. After ablation, the anti-Ucn3 also marked bihormonal cells, confirming our RNAseq data (Figure 4I). Based on these new transcriptomic datasets, we defined the genes selectively enriched (>4-fold) in *sst1.1* δ-cells versus the other endocrine cell types already available (*sst2* δ-, β- and α-) and identified 152 specific GFP^high^*/sst1.1* δ-cell makers, among which *bdnf, cdh10a, sox11b* and *dkk3b* (Figure 4-Source Data 3). An updated list of 60 markers enriched in β-cells versus *sst1.1* δ-cells, α- and *sst2* δ-cells could also be defined. Our RNAseq data also revealed that *dkk3b* and *ucn3l*, previously attributed to β-cells, were even more enriched in GFP^high^*/sst1.1* δ-cells.

Top GO terms overrepresented in GFP^low^/*sst2* δ-cells (Figure 4J and Figure 4-Source Data 4) were related to neuron differentiation *(isl1, tbx2b*, *map1ab*…), adhesion (*cdh2, alcamb, nrcama…)*, especially focal adhesions and ECM-receptor interaction (*tnc, itga1, col6a1…*), plasma membrane projection (*vegfaa, ngfa, sema4c…*), Wnt signalling (*wnt9a, tcf7l1b, ccnd1…*), and p53 signalling with the *rprma, sesn1, apaf* genes that were found also enriched in the *sst2* δ-cell dataset from (Tarifeño-Saldivia et al., 2017).

Top most significant GO terms and pathways in GFP^high^/*sst1.1* δ-cells (Figure 4J and Figure 4-Source Data 5) identified regulation of communication and signalling (*dkk3b, pde3a, chrm4a*…), adhesion (*nlgn1, ncam1b, cdh10a, dlg1, celsr1a, ctnnb1, cadm1a/b …*), regulation of localization (*chga, sema3ga/b, cacna1b*…) and metabolism of carbohydrates (*gck, g6pcb, slc2a2, hk2…).* In line with this metabolic signature, the key regulator of energy metabolism in glucose- and Insulin-sensitive cells, *foxo1a/b*, was also overexpressed in GFP^high^*/sst1.1* δ-cells. Furthermore, enriched expression in GFP^high^*/sst1.1* δ-cells of *gck*, the limiting glucose sensor in β-cells and *slc2a2/*GLUT2, the low affinity glucose transporter utilized by β-cells for glucose intake, denotes competence for glucose-responsive hormone secretion. Several proprotein convertases with hydrolase activity important in the secretory pathway, *pcsk1, pcsk2, dpp6b*, a β-cell gene, were also identified in GFP^high^*/sst1.1* δ-cells.

Overall, GFP^high^*/sst1.1* and *sst1.1*GFP^low^/*sst2* δ-cells express specific repertoires of adhesion molecules and surface receptors reflecting different intracellular signalling activities and cellular phenotypes. Importantly, these data unveil that GFP^high^*/sst1.1* δ-cells represent a distinct δ-cell population possessing the basic features of β-cells and sensors to integrate INS signalling and glucose metabolism and carry hormone secretory activity; all these features make them ideal candidates to adapt to the loss of β-cells, induce *ins* expression and convert to bihormonal cells.

### Bihormonal cells derive from sst1.1 δ−cells that induce ins expression after β-cell ablation

To explore the relationship between GFP^high^*/sst1.1* δ-cells and bihormonal cells, we quantified the GFPhigh/*sst1.1* δ-cell GFP population during regeneration. At 3 and 20 days after β-cell ablation, GFP^high^*/sst1.1* (ins:Cherry-) δ-cells were markedly depleted (Figure 5A), and this reduction was paralleled by the induction of bihormonal cells observed in Figure 1, therefore strongly arguing that GFP^high^*/sst1.1* δ-cells switched to GFP^high^*/sst1.1+ ins:*Cherry+ cells. Then, we followed the appearance of bihormonal cells by *in vivo* time lapse imaging of the main islet following ablation in 3 dpf *Tg(sst1.1:eGFP); Tg(ins:NTR*-P2A-mCherry)* larvae. Figure 5B-B’ show Cherry fluorescence progressively appearing in pre-existing *sst1.1:eGFP* cells presenting strong GFP fluorescence, presumably GFP^high^*/sst1.1* δ-cells. This demonstrates that GFP^high^*/sst1.1* δ-cells directly convert into bihormonal cells.

**Figure 5.**
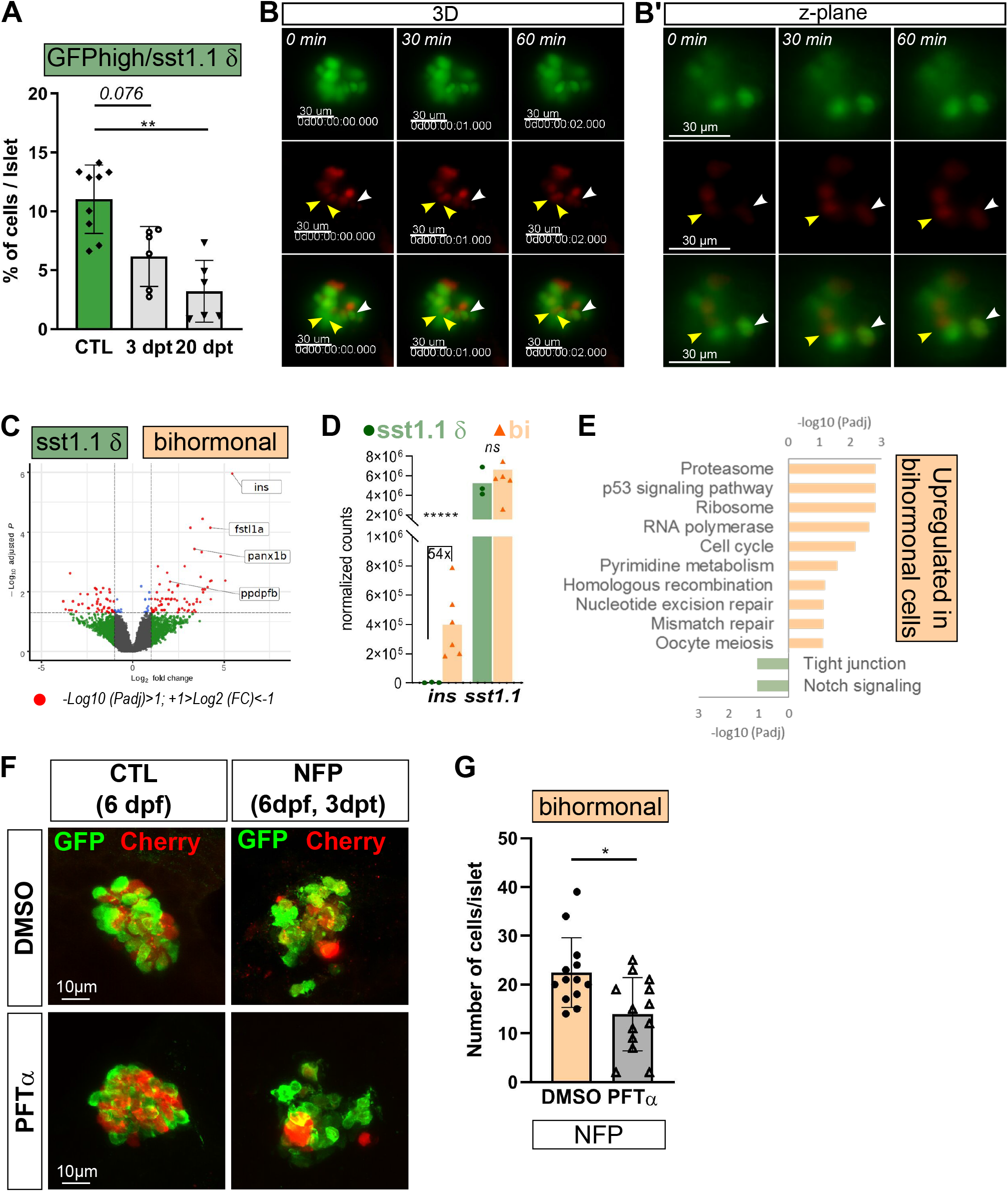
*sst1.1* δ-cells convert to Sst1.1+ Ins+ bihormonal cells after β-cell destruction and require p53 activity. A) Quantification by flow cytometry of GFP^high^/*sst1.1* δ-cells before ablation (CTL) and at 3 and 20 dpt depletion of GFP^high^/*sst1.1* δ-cells. Cells were isolated from dissected main islets of adult *Tg(sst1.1:eGFP); Tg(ins:NTR*-P2A-mCherry)*. Mean ± SD; Kruskal-Wallis test; \*\**P<0.005* B) *In vivo* time lapse of the main islet of a 4 dpf *Tg(sst1.1:eGFP); Tg(ins:NTR*-P2A- mCherry)* larva following β-cell ablation from 3 to 4 dpf. 3D representation (B) and one z-plane (B’) of the same islet are shown. The arrowheads point at two GFP+ cells (green) that start to express ins:mCherry (red) fluorescence between *t0* and *t1* (visible in the same z-plane). One of the sst1.1+ presents strong GFP fluorescence (white arrowhead). Images were acquired every 30 min starting from 4 dpf (96 hpf). C) Volcano plot showing the significant DE genes over- or under-expressed in 20 dpt bihormonal cells versus CTL GFP^high^/*sst1.1* δ-cells (FC>2<, Padj<0.05). The full list of significant DE genes calculated by DESeq is provided in Figure 5-Source Data 1. D) Expression in normalized counts of the *sst1.1* and *ins* genes in CTL GFP^high^/*sst1.1* δ-cells and bihormonal cells (bi). Padj are calculated by DESeq. *ns:* no significant DE between the two conditions, *****<0.000005. E) Top significant KEGG pathways identified among the genes upregulated (in orange) and downregulated (in green) in bihormonal cells compared to CTL GFP^high^/*sst1.1* δ-cells. The list of GO terms below FDR 0.25 is given in Figure 5-Source Data 2 and 3. F) Confocal images of the main islet. β-cell ablation was performed in *Tg(sst1.1:eGFP); Tg(ins:NTR*-P2A-mCherry)* by NFP treatment from 3 to 4 dpf embryos then larvae were exposed to the p53 inhibitor pifithrin α (PFTα) from 4 to 6 dpf. Representative images are shown for each condition (3D projections). G) Quantification of bihormonal and sst1.1:GFP+ cells (GFP only) after ablation (NFP) and treatment with DMSO (CTL) or PFTα based on the confocal images. Mean ± SD; Unpaired Mann-Whitney test; \**P<0.05, ***P<0.0005*.

To further assess the similitudes between these two cell types, we compared their transcriptome. Quite few DE genes were identified with 293 over- and 180 underexpressed genes in bihormonal cells versus GFP^high^*/sst1.1* δ-cells (FC 2-fold, Padj <0.05) (Figure 5C and Figure 5-Source Data 1). *ins* was the top overexpressed gene in bihormonal cells (54-fold) (Figure 5D). Among the 293 overexpressed genes in bihormonal cells, 9 were β-cell markers, such as *ins*, *fstl1a* and *panx1b*, and 8 genes out of the 180 underexpressed ones were GFP^high^*/sst1.1* δ-cell markers. Both *sst1.1* and *hhex* were maintained, underscoring that bihormonal cells retain most of the *sst1.1* δ-cell identity. The most significant enriched pathways amongst the overexpressed genes in bihormonal cells were “ribosome”, “proteasome”, “p53 signaling pathway” and “cell cycle” with several genes involved in cell cycle arrest (Figure 5E and Figure 5-Source Data 2) suggesting p53-mediated cell cycle arrest and protein homeostasis response. The only significant downregulated pathways were linked to adhesion (tight junction and Notch signalling) (Figure 5-Source Data 3).

Overall, these data highlight a remarkable resemblance in terms of cell identity between GFP^high^*/sst1.1* δ-cells and bihormonal cells. The “pro β-“ *sst1.1* δ-cells switch on *ins* expression through direct conversion via mechanisms possibly involving the p53 pathway and reduced adhesion and cell-cell contacts.

### Inhibition of the p53 pathway compromises formation of bihormonal cells

The role of the p53 pathway in bihormonal cell formation was next assessed. β-cell ablation was performed at 3 dpf then followed by treatment with the p53 inhibitor Pifithrin-α (PFTα) from 4 to 6 dpf or DMSO (Figure 5F-G). While PFTα did not affect islet cells in CTL larvae, quantification of double GFP+ Cherry+ cells after β-cell ablation revealed fewer bihormonal cells in larvae exposed to the inhibitor (Figure 5G). These results strongly suggest that p53 promotes the formation of bihormonal cells.

### Bihormonal cells persist long after β-cell ablation

Next, we questioned the persistence of bihormonal cells long after ablation and analysed the main islet from *Tg(sst1.1:eGFP); Tg(ins:NTR*-P2A-mCherry)* adult fish 4 months after ablation. Surprisingly, most Ins+ cells still co-expressed GFP as well as high levels of Ucn3 at this stage (Figure 6A), similarly to 20 dpt bihormonal cells. Bihormonal cells still constituted the vast majority of *ins*-expressing cells in the main islet compared to mCherry+ (GFP-) β-cells (Figure 6B-C) showing that they do not represent a transient intermediary population that would ultimately resolve into *ins*-only β-cells.

**Figure 6.**
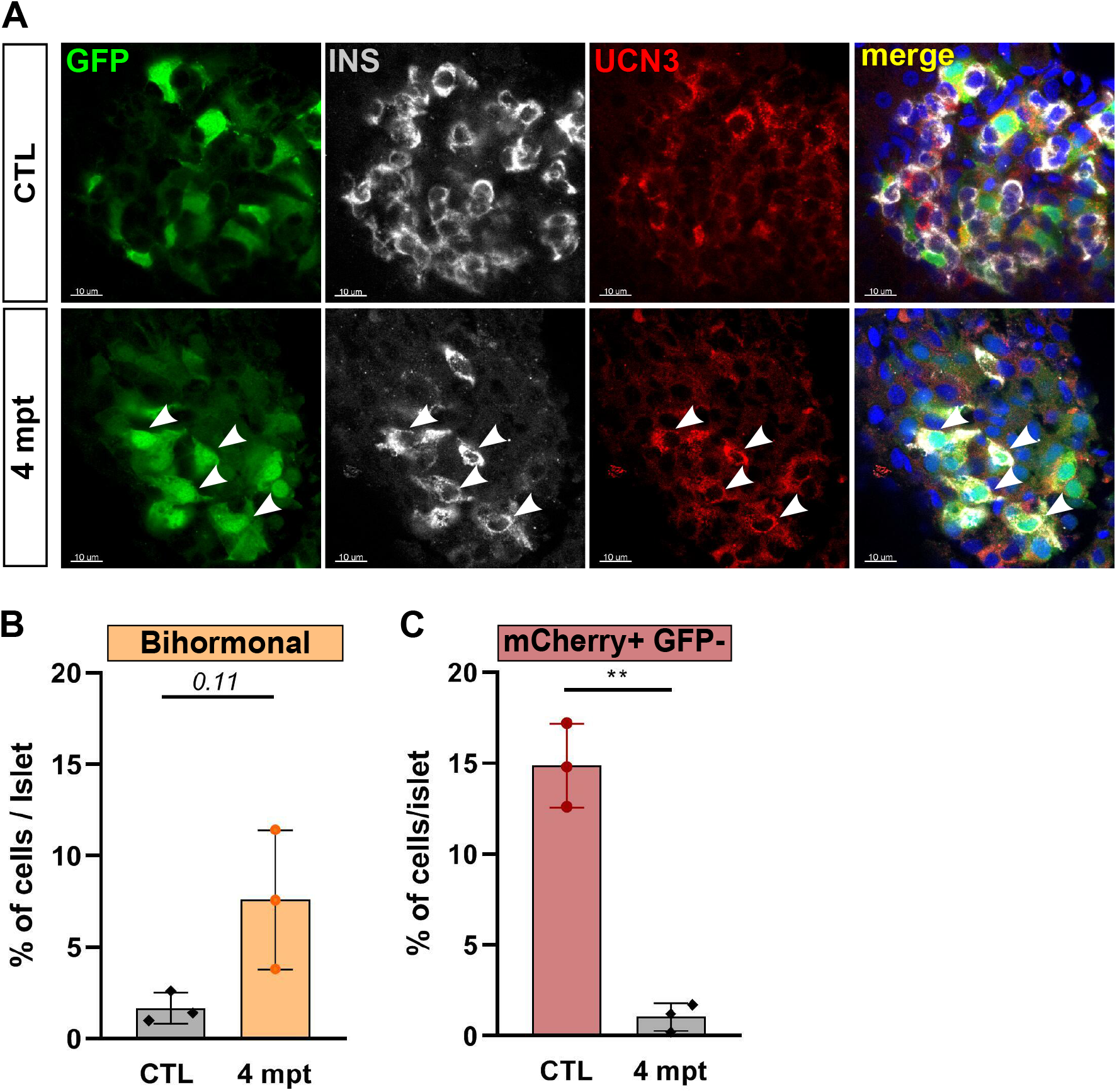
Protracted bihormonal cells 4 months after β-cell ablation. A) Whole mount immunodetection of UCN3 (red), GFP (green), INS (grey) in the main islet of *Tg(sst1.1:eGFP); Tg(ins:NTR*-P2A-mCherry)* revealing persistent bihormonal GFP+ INS+ cells still 4 months after ablation. These cells still also express Ucn3 (white arrowheads). B-C) Quantification by flow cytometry of mCherry+GFP+ bihormonal cells (B) and mCherry+GFP- β-cells (C) populations in CTL and 4 months after ablation by flow cytometry. Means ± SD; Unpaired t-test with Welch’s correction. \**\*P<0.005*

### Bihormonal cells account for the main ins-expressing cells responsible for blood glucose control

Bihormonal cell express the basic machinery for glucose-induced Ins secretion and the glycemia of regenerated fish is normalized. This strongly suggests that these cells contribute to blood glucose control. To exclude the possibility that glycemia is regulated by another population of regenerated β-cells outside the main islet, we analysed secondary islets. In addition to the large main islet located in the pancreatic head, zebrafish possess smaller secondary islets which are scattered in the pancreatic tail. Immunofluorescence revealed that, like the main islet, CTL secondary islets do not display significant overlap between β-cells (Cherry+) and *sst1.1*:GFP+ cells. Furthermore, similarly to main islets, 20 dpt secondary islets harboured many bihormonal cells and very scarce Cherry+ (GFP-) β-cells (Figure 7A). Thus, as bihormonal cells constitute the predominant *ins*-expressing cell population throughout the whole pancreas, it can be concluded that they are responsible for the normalization of glycemia after ablation and regeneration.

**Figure 7.**
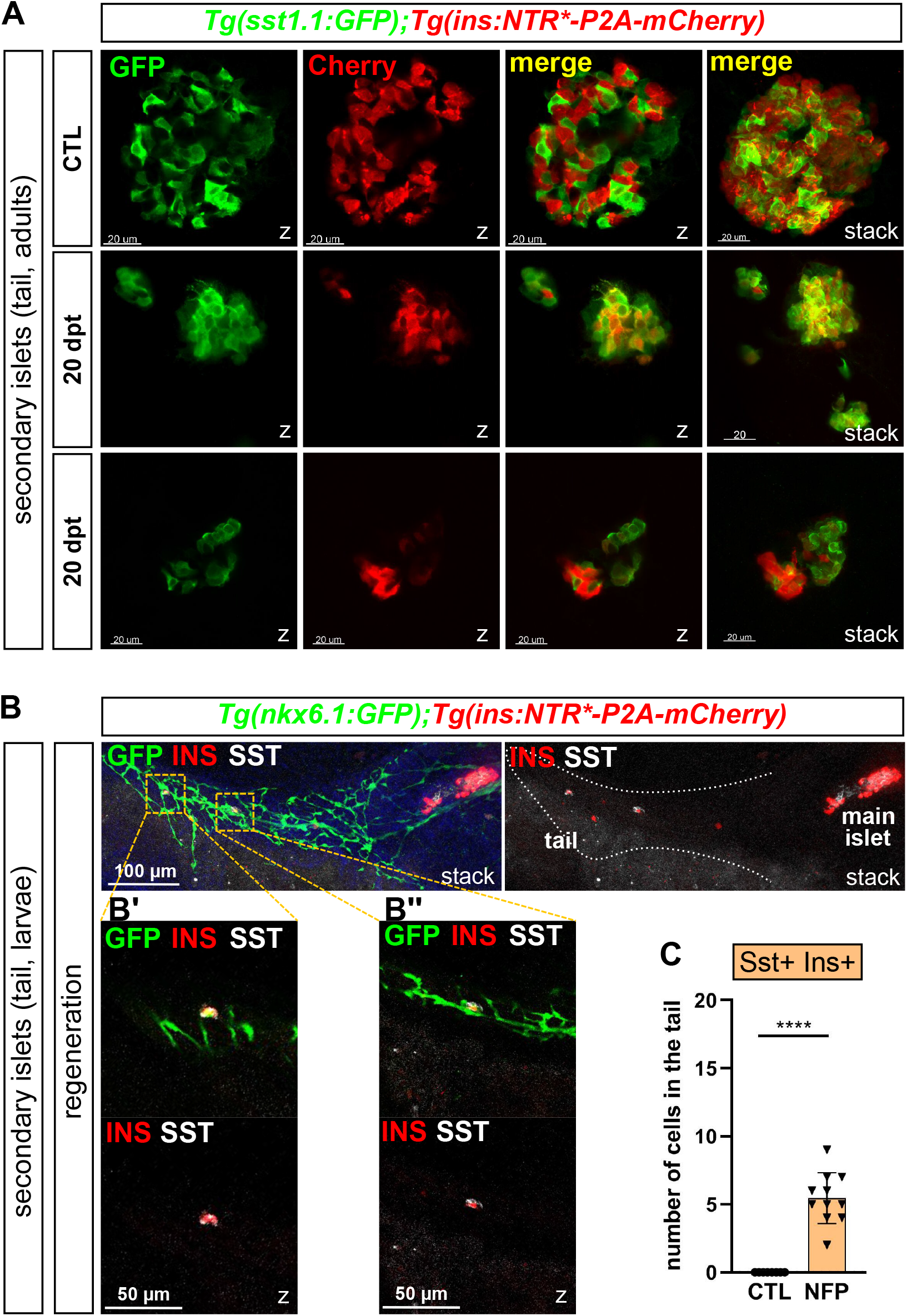
Bihormonal cells in secondary islets and from ductal origin. A) Immundofluorescence (GFP and mCherry) on *Tg(sst1.1:eGFP); Tg(ins:NTR*-P2A-mCherry)* showing representative CTL and 20 dpt secondary islets present in the pancreatic tail of adult zebrafish showing that most *ins*-expressing cells (Cherry+) are still bihormonal (GFP+), like in the main islet. Two independent samples are shown at 20 dpt. Cells appear in yellow due to overlapping GFP and mCherry staining. Z-planes and 3D projections (stack) are shown. B) Representative confocal images showing duct-derived Ins+ cells in the tail in *Tg(nkx6.1:eGFP)*; *Tg(ins:NTR*-P2A-mCherry)* larvae after treatment with NFP from 3 to 4 dpf. Larvae were analysed at 17 dpf for GFP (duct cells), Ins (red) and Sst (white) immunolabelling, here shown in condition of regeneration. Stacks represent 3D projections of the whole pancreas and close-ups of individual bihormonal cells in the tail in B’ and B’’ are shown as z-planes. The pancreatic tail is delineated by a white dashed line. C) Quantification based on confocal images. CTL larvae do not show any bihormonal cells while post-ablated larvae display Ins+ Sst+ cells. Mann-Whitney test, *\*\*\*\*P<0.0001*

### Bihormonal cells also arise from pancreatic ducts

In zebrafish, the secondary islets originate from pancreatic ducts (Parsons et al., 2009; Wang, Rovira, Yusuff, & Parsons, 2011) and β-cell regeneration can also occur from duct cells in the adult zebrafish (Delaspre et al., 2015; Ghaye et al., 2015). The striking observation that the vast majority of new *ins*-expressing cells are bihormonal in the entire pancreas raised the hypothesis that duct-derived Ins+ cells also express Sst1.1. To explore this possibility, we used larvae, a well-established model to study β-cell regeneration from the ducts (Ninov et al., 2013)(Ninov, Borius, & Stainier, 2012), and monitored Ins and Sst expression along the main duct in the tail using *Tg(nkx6.1:eGFP); Tg(ins:NTR*-P2A-mCherry)* (Ghaye et al., 2015) (Figure 7B). Ablation was performed at 3 dpf when no differentiated β- and δ-cells are yet present in the tail, which means that potential new Ins+ Sst1.1+ cells following ablation originate from the ducts. Similar to the islets, bihormonal cells were induced in the ductal GFP+ domain in the tail of regenerating larvae (Figure 7B-C), indicating that duct cells can give rise to bihormonal cells and contribute to the overall bihormonal cell mass.

## Discussion

Pancreatic endocrine cell plasticity has emerged as an important cellular adaptive behaviour in response to β-cell stress and death in human and in mammalian diabetic models. Here we show that, in zebrafish, the plasticity of highly differentiated endocrine cells is a major contributor to β-cell regeneration, but, in contrast to mammals, a specific subpopulation of δ-cells with a “pro-β-cell” potential confers very efficient and fast adaptation to β-cell loss all along zebrafish life.

The rapid formation of so many INS+ SST1.1+ bihormonal cells following β-cell ablation strongly suggests that they directly arise from a pre-existing islet cell type with close features. GFP^high^*/sst1.1* δ-cells represent ideal candidates for this adaptation. First, pre-existing GFP^high^*/sst1.1* δ-cells already express about half the previously defined β-cell genes such as *pdx1, ucn3l* and *slc2a2* genes and their transcriptomic profile is very close to bihormonal cells. Second, δ- and β-cells use the same glucose-stimulated and calcium-dependent mechanisms for hormone secretion ((Denwood et al., 2019; Rorsman & Huising, 2018) for review). Indeed, zebrafish GFP^high^*/sst1.1* δ-cells express the basic molecular machinery for glucose-sensing and glucose-dependent stimulation of insulin secretion and blood glucose control. Third, the appearance of bihormonal cells concurs with depletion of the GFP^high^*/sst1.1* δ-cell mass after regeneration. Fourth, we excluded pre-existing β-cells as the source of bihormonal cells and the other δ-cell subpopulation, the *sst2* δ-cells, have previously been ruled out as potential origin of β-cell regeneration in zebrafish (Ye, Robertson, Hesselson, Stainier, & Anderson, 2015). Finally, *in vivo* imaging revealed that GFP^high^*/sst1.1* δ-cells indeed turn on *ins:*mCherry expression. All these observations bolster that *sst1.1* δ-cells are δ-cells with β-cell features enabling them to rapidly reprogram to bihormonal cells by activating *ins* expression and engender functional surrogate β-cells that can compensate for the loss of β-cells.

The phenomenon of age-independent direct conversion of δ-cells to β-cells in zebrafish contrasts with the situation in mouse where δ-cells can only transiently regenerate β-cells at neonatal stages through a process involving dedifferentiation, proliferation and activation of *Ins* (Chera et al., 2014). On the other hand, the phenomenon of bihormonal cell formation in zebrafish shares similitudes with the direct conversion operated from α-cells in adult mice which transiently produce GCG+ INS+ bihormonal cells (Thorel et al., 2010). Nevertheless, there are also striking differences such as a very fast and efficient process in zebrafish as opposed to the slow and limited conversion of mouse α-cells (Thorel et al., 2010). While only a small fraction of α-cell convert in mouse, zebrafish *sst1.1* δ-cells seem to represent a reservoir of “pro-β” cells ready for massive conversion towards bihormonal cells at any stage, underlining the potent plasticity of zebrafish *sst1.1* δ-cells. The plasticity of murine α-cells has been shown to be restricted by the synergistic action of Ins and of several, many of them unknown, signals which precludes their massive conversion. Interestingly, murine adult δ-cells contribute to restrict β-cell fate in α-cells (Cigliola et al., 2018). Of note, in zebrafish too, α-cells have been shown to transdifferentiate into β-cells via activation of Pdx1 (Lu et al., 2016; Ye et al., 2015) showing that complex cross-talks between several islet cell populations are involved in the adaptation to the loss of β-cells throughout species. In conclusion, zebrafish *sst1.1* δ-cells are already predisposed to convert to Ins+ cells at basal state; their plasticity and conversion into Ins-expressing cells represent an ancestral means of adaptation to the loss of β-cells that is fully expressed in zebrafish but has become limited in mammals.

The efficient conversion of *sst1.1* δ-cells to *ins*-expressing cells and the intrinsic competence of *sst1.1* δ-cells to induce *ins* might be conferred by the expression of the Pdx1 transcription factor. However, *pdx1* expression alone is obviously not sufficient to guarantee *ins* expression, and other mechanisms consequent to β-cell loss must operate in synergy, such as epigenetic regulations. For example, Pdx1 transcriptional activity has been shown to be controlled by a glucose-induced switch from Pdx1/BRM repressor to Pdx1/BRG1 activator complexes allowing maximal expression of *Ins* and *Ucn3* (McKenna, Guo, Reynolds, Hara, & Stein, 2015). In agreement with this, *smarca2* (BRM) was downregulated in bihormonal cells (0.54x), possibly resulting in a shift towards high level of the Brg1/Pdx1 activator complex. Further modifications of the epigenetic chromatin state via histone methylation and acetylation at the level of the *ins* locus could be involved in the consolidation of the bihormonal phenotype. The study of *sst1.1* δ-cells and of the mechanisms that reinforce their β-cell identity upon β-cell loss should help restore a pool of functional INS-producing cells in diabetic patients.

Although zebrafish bihormonal cells express the essential β-cell determinants *ins* and *pdx1*, they totally lack two other key β-cell genes, *nkx6.2* and *mnx1*, leading to the surprising conclusion that none of these two genes is pivotal for *ins* expression despite their importance for β-cell specification during development (Binot et al., 2010; Gokhan Dalgin et al., 2011; Henseleit et al., 2005; Schaffer et al., 2013). Our results are rather more consistent with another important role of these genes in repressing non-beta endocrine lineage programs, particularly delta, like in their murine orthologues (Gökhan Dalgin & Prince, 2012; Pan et al., 2015; Schaffer et al., 2013). In addition, zebrafish bihormonal cells have lower levels of *nkx2.2a* whose orthologue in mammals also represses δ-cell features (Gutiérrez et al., 2017). On the other hand, *nkx6.1*, totally absent in GFP^high^*/sst1.1* δ-cells, is detected in bihormonal cells (Figure 5-Source Data1) and might partially compensate for the absence of the β-cell marker *nkx6.2* in conferring β-cell identity. Together, low or absent expression of these genes likely underlines the mixed delta-beta phenotype.

The present study provides an in-depth characterization of the zebrafish *sst1.1+* δ-cell subpopulation and corroborates recent reports of the existence of two clusters of δ*-*cells by single cell RNAseq (Spanjaard et al., 2018), one expressing *sst2/sst1.2* and the other *sst1.1*. Preferential expression of the *sst1.1* δ-cell markers we identified here such as *bdnf, cdh10a, sox11b* and *dkk3b* was confirmed in the *sst1.1+* cluster by a reanalysis of these single cell RNAseq data (not shown). Distinct δ-cell populations differentially expressing three *sst* genes highlights a phenomenon of sub-functionalization. In mammals, SST peptides are generated by cleavage of the SST(1-28) pro-hormone. The predominant pancreatic form, SST(1-14), is produced via cleavage by Pcsk2 which is highly enriched in GFP^high^*/sst1.1* δ-cells. The Sst(1-14) peptide predicted to be produced from the zebrafish Sst1.1 prohormone is 100% conserved with human SST(1-14). The Sst(1-14) peptide deriving from Sst1.2 has 86% identity with human SST(1-14) peptide while the one deriving from Sst2 is the most divergent with 64% identity with SST(1-14) (Devos et al., 2002). All this strongly suggests that Sst1.1 (*sst1.1* δ-cells) and Sst1.2 (*sst2* δ-cells) are likely the functional orthologues of mammalian SST. This raises the hypothesis that *sst2* δ-cells may maintain δ-cell function following conversion of *sst1.1* δ-cells.

Also, GFP^low^*/sst2* δ-cells express *chrm3a* known to stimulate SST secretion in mammalian δ-cells (Rorsman & Huising, 2018) while *sst1.1* δ-cells expressed higher level of the inhibitory *chrm4a*. Thus the transcriptomic profiles of GFP^high^*/sst1.1* δ-cells and GFP^low^*/sst2* δ-cells confirm the sub-functionalization in terms of secretory activities in addition to the competence to express *insulin*. Of note, human δ-cells, like zebrafish GFP^high^*/sst1.1* δ-cells, express *PDX1.* Although the occurrence of SST+ INS+ cells in mammals remains largely undocumented, single cell RNAseq data suggest the existence of putative SST+ INS+ bihormonal cells in adult human islets (http://sandberg.cmb.ki.se/pancreas/ (Segerstolpe et al., 2016)) and in NOD murine islets (Thompson et al., 2019). It will be interesting to determine whether they derive from a competent subpopulation of δ-cells that could be stimulated to help restore glucose homeostasis in diabetic patients.

With their low *ins* expression and high *glucokinase* (*gck)* expression, bihormonal cells are reminiscent of “hub”, or “leader” cells, a subpopulation of β-cells identified in mammals (Johnston et al., 2016) and recently in zebrafish (Salem et al., 2019). These cells rapidly respond to glucose and play the role of pacemakers that connect the other β-cells to coordinate their activity. Interestingly, zebrafish bihormonal cells express the “vascular muscle contraction” signature and the *cacna1h* and *cacna1i* genes which are also enriched in neurons that have pacemaker activity. Further similitude with neurons is underscored by expression of the neuronal marker *elavl3* (HuC) in GFP^high^*/sst1.1* δ-cell (Figure 4-Source Data 1) and bihormonal cells (Figure 3-Source Data 1). Indeed, *elavl3* promoter activity has been previously observed in some undefined endocrine cells of the pancreatic islet (Yang, Kawakami, & Stainier, 2018). From our study, it is tempting to hypothesize that zebrafish hub cells and bihormonal cells share a common lineage with GFP^high^*/sst1.1* δ-cells.

Although most of this study was focused on the main islet, we also report that bihormonal cells constitute the majority of *ins*-expressing cells also in the pancreatic tail which supports conversion from *sst1.1* δ-cells in the secondary islets. Duct cells can also give rise to regenerated β-cells (Delaspre et al., 2015; Ghaye et al., 2015). We show that even duct-derived Ins+ cells can be bihormonal, which further contributes to the prevalent bihormonal phenotype versus *bona fide* β-cells after regeneration. Whether duct-derived bihormonal cells differentiate via a *sst1.1*-only transitional state remains to be determined.

Bihormonal cells display signatures associated with p53 signalling, proteostasis and cell cycle arrest. In addition, inhibition of p53 activity reduces the number of bihormonal cells. In response to various stress such as ROS and metabolic stress, p53 promotes cell cycle arrest and responses such as metabolic adaptation and DNA repair, or apoptosis when the pro-survival mechanisms are overwhelmed (Hafner, Bulyk, Jambhekar, & Lahav, 2019). Collectively, these results suggest that bihormonal cell formation depends on the protective activity of p53 as a result of adaptation to the loss of β-cells. Of note, bihormonal cells were observed using two different ablation models, NTR/NFP and DTA, both systems involving acute β-cell death, loss of Ins signalling, islet niche disruption, release of genotoxic ROS and hyperglycemia. The role of these signals remain unknown whilst ROS, hyperglycemia and loss of Ins signalling are not sufficient to drive formation of Ins+ Sst1.1+ bihormonal cells (our data not shown), indicating that this process requires additional signals resulting from β-cell destruction. As bihormonal cells not only appear in the main islet and in secondary islets but also in the pancreatic ducts, it is tempting to hypothesize that their formation in these different compartments share common mechanisms.

Normal glycemia is recovered after 20 days despite the very low abundance of genuine β-cells. They also constitute a stable and prevalent population long after ablation. Given that this population of bihormonal cells account for the vast majority of Ins+ producing cells throughout the pancreas, we conclude that they are the functional unit involved in normoglycemia in regenerated fish. Moreover, their capacity to regulate blood glucose levels is corroborated by their transcriptomic profile showing that they possess the energy metabolism and the machinery required for glucose responsiveness and insulin secretion.

Efficient neogenesis of functional Ins-producing cells *in situ* would have a great impact in diabetes in humans. These findings illustrate the importance of understanding the similitudes between zebrafish *sst1.1* δ-cells and β-cells and the mechanisms underlying the conversion of *sst1.1* δ-cells to Ins+ bihormonal cell for translational perspectives.

## Materials and Methods

### Zebrafish husbandry and generation of the Tg(sst1.1:eGFP)^ulg054^ zebrafish line

Zebrafish wild-type AB were used in all the experiments. *TgBAC(nkx6.1:eGFP)^ulg004^* (Ghaye et al., 2015) and *Tg(ins:NTR*-P2A-mCherry)^ulg034^* (Bergemann et al., 2018) were used. Zebrafish were raised in standard conditions at 28°C. All experiments were carried out in compliance with the European Union and Belgian law and with the approval of the ULiège Ethical Committee for experiments with laboratory animals (approval numbers 14-1662, 16-1872; 19-2083).

To generate the *Tg(sst1.1:eGFP)^ulg054^* zebrafish line, the *sst1.1:eGFP* transgene has been generated by cloning a 770 pb PCR fragment containing the *sst1.1* regulatory regions just upstream the ATG of the *sst1.1* ORF (ENSDARG00000040799.4) amplified with primers IM217 (reverse: 5’-TTTTATTAAAGTGTTTATTTGGTCTCAGAG-3’) and IM256 (forward: 5’-AAGAGCACTTCAGATGTCTTCCC-3’) into the Gateway vector pCR8/GW/TOPO. The promoter was assembled by LR recombination with p5E-MCS and p3E-eGFP into pDestTol2p2A from the Tol2kit (Kwan et al., 2007). *Tg(sst1.1:eGFP)^ulg054^* fish have been generated using the Tol2 mediated transgenesis (Kawakami, 2007). Adult *Tg(sst1.1:eGFP)^ulg054^* fish (abbreviated *Tg(sst1.1:eGFP))* were crossed with *Tg(ins:NTR*- P2A-mCherry)^ulg034^* to generate a double transgenic line. The insbglob:loxP-mCherry-nls-loxP-DTA construct was created by cloning a loxP-mCherry-nls loxP cassette downstream of the *ins* promoter beta-globin intron (Ninov et al., 2013). Subsequently, a DTA gene was cloned downstream of the last loxP site via ligation independent cloning (InFusion, Clontech). The *Tg(ins.bglob:loxP-NLS-mCherry-loxP-DTA)^bns525^* line (abbreviated *Tg(ins:lox-mCherry-lox-DTA*)) was generated using the Tol2 system (Kawakami, 2007). *The Tg(ins:CRE-ERT2*) has been generated by LR recombination combining p5E-MCS (Kwan et al., 2007), pME-ins and p3E-CRE^ERT2^ vectors into pDestTol2p2A from the Tol2kit. pME-ins was obtained by cloning into the pCR8/GW/TOPO a PCR fragment of 897 pb using the primers O097 (GTATCTATAGTTGAACATGAAAGCAT) et O098 (GGTCACACTGACACAAACAC ACA) and which contains 744 bp of the insulin promoter, the exon 1 (47 bp), the intron 1 (99 bp) and the 7 bp of exon 2 just upstream of the ATG. p3E-CRE^ERT2^ was obtained by BP cloning into the pDONRP2R-P3 the 2200bp PCR fragment using the primers O99 GGGGACAGCTTTCTTGTACAAAGTGG CTGCTAACCATGTTCATGCCTTC and O100 GGGGACAACTTTGTATAATAAAGTTGTCAAGCTGTGGCAGGGAAACCC and as template the pCRE^ERT2^ kindly received from P. Chambon(Feil, Wagner, Metzger, & Chambon, 1997).

### β-cell ablation

Nifurpirinol (NFP) (32439, Sigma-Aldrich) stock solution was dissolved at 2.5 mM in DMSO. 4-Hydroxytamoxifen (4-OHT, H7904, Sigma-Aldrich) was dissolved in DMSO as a concentrated solution of 10 mM and kept as single-use aliquots at –80 °C. β-cell ablation in *Tg(sst1.1:eGFP); Tg(ins:NTR*-P2A-mCherry)* larvae was induced by treatment with 4 µM NFP in E3 egg water. Adult fish were treated in fish water with 2.5 µM NFP. Control treatments consisted of E3 containing 0.16% DMSO. Larvae and adults were treated for 18 hours in the dark. To induce β-cell ablation with *Tg(ins:lox-mCherry-lox-DTA); Tg(ins:CRE-ERT2)* line, larvae were treated with 5 µM 4-OHT 2x 2 hours in the dark.

### Blood glucose measurements

Adult fish were fasted for 24 hours then euthanized with tricaine and the glycemia was immediately measured using the Accu-Chek Aviva glucometer (Roche Diagnostics) with blood collected at the tail.

### Immunodetection of paraffin sections

Samples were fixed and processed for immunofluorescence as previously described (Ghaye et al., 2015).

### Whole mount immunodetection

Larvae were euthanized in tricaine and fixed in 4% PFA at 4 °C for 24 hrs before IHC. After depigmentation with 3% H2O2/1% KOH during 15 min, larvae were permeabilised 30 min in PBS/0.5% Triton X-100 and incubated for two hours in blocking buffer (4% goat serum/1% BSA/PBS/0.1% Triton X-100). Primary and secondary antibodies were incubated at 4 °C overnight. Adult fish (6-10 months) were euthanized and fixed for 48 hrs. Digestive tracts were dissected, dehydrated and stored in 100% methanol at −20 °C. Before IHC, the samples were permeabilised in methanol at room temperature for 30 min, placed 1 hr at −80 °C then back at room temperature. After rehydration in PBS/0.05% Triton X-100, depigmentation was performed for 15 min followed by incubation in blocking buffer containing 4% goat serum /1% BSA/PBS/0.01% Triton X-100. The primary antibodies were incubated for 48 hrs on adult samples and overnight on larvae, followed by overnight incubation with the secondary antibodies overnight at 4 °C. Primary antibodies: Anti-Insulin (guinea pig, 1:500, Dako A0564), Living Colors Polyclonal anti-mCherry/dsRed (rabbit, 1:500, Clontech 632496), Living Colors Polyclonal anti-Pan-RCFP (rabbit, 1:500, Clontech 632475), anti-GFP (chicken, 1:1000, Aves lab GFP-1020), anti-Somatostatin (rat, 1:300, Invitrogen MA5-16987), anti-Glucagon (mouse, 1:300, Sigma G2654), anti-Urocortin 3 (rabbit, 1:300, Phoenix Pharmaceuticals H-019-29), anti-Pdx1 (guinea pig, 1:200, kind gift from Chris Wright, Vanderbilt University). Secondary antibodies: Alexa Fluor-488, -568, -633 (goat, 1:750, Molecular Probes).

### Whole mount in situ hybridization on embryos

The *sst1.1* probe was synthesised with SP6 and the *sst2* probe with T7(Devos et al., 2002). Fluorescent in situ hybridization were performed as described in(Tarifeño-Saldivia et al., 2017) on 3 days post fertilization embryos (dpf). The antisense RNA probes were revealed using tyramide-Cy3 followed by immunodetection of GFP. Images of immunodetection and in *situ* hybridization were acquired with a Leica SP5 confocal microscope, and processed with Imaris 9.5 (Bitplane) for visualization.

### In vivo imaging

*In vivo* imaging was performed with a Lightsheet Zeiss Z1 microscope using a 20x water immersion objective and 488nm and 561nm lasers. *Tg(sst1.1:eGFP); Tg(ins:NTR*-P2A-mCherry)* larvae were treated from 1 dpf with 1-phenyl 2-thiourea (0.003% (w:v)) to inhibit pigment synthesis. After ablation with NFP from 3 to 4 dpf, larvae were anesthetized, embedded in 0.25% low melting agarose containing and mounted into FEP capillaries. Images were acquired every 30 min and were maintained during the whole experiment at 28° and with 100 ml/L tricaine. Images were converted with Imaris 9.5 (Bitplane) for visualization.

### Flow cytometry and FACS

The zebrafish pancreas contains one main big islet in the head and several smaller secondary islets in the tail. The main islets from 2-4 pancreata of *Tg(sst1.1:eGFP); Tg(ins:NTR*-P2A-mCherry)* adult fish (6–10 months old, males and females) were dissected under epifluorescence to eliminate a maximum of non-fluorescent surrounding exocrine tissue, collected and washed in HBSS without Ca^2+^/Mg^2+^. Live cell dissociation was performed in Tryple Select 1x solution (GIBCO) supplemented with 100 U/mL collagenase IV (Life Technologies 17104-019) and 40 µg/mL proteinase K (Invitrogen, 25530031) for 10 min at 28 °C, and stopped with 15% FBS. The GFP+ cells, mCherry+ cell and double GFP+ mCherry+ cells were selected according to gates as shown in Figure 1-figure supplement 2 (dashed lines) on FACS Aria III and sorted under purity mode and after exclusion of the doublets. The purity of the sorted cells was confirmed by epifluorescence microscopy (∼95 %). Cells (about 1000-5000/fish depending on the cell type) were immediately lysed with 0.5% Triton X-100 containing 2U/µl RNAse inhibitor and stored at –80 °C.

### Cell quantification in adults by flow cytometry

The percentage of mCherry+, GFP+ and double mCherry+ GFP+ fluorescent cells in the dissociated islets was inferred from flow cytometry experiments in each quadrant delimiting negative and positive fluorescence. FACS plots were generated by FlowJo 10.6.2.

### mRNA sequencing of FACSed cells and bioinformatic analyses

cDNAs were prepared from lysed cells according to SMART-Seq2.0 (Picelli et al., 2014) for low input RNA sequencing and libraries were prepared with Nextera® DNA Library kit (Illumina). Independent biological replicates of each cell type sequenced using Illumina HiSeq2500 and obtained ∼20 million 75 bp single-end reads (7 replicates for β-cells, 6 for 20 dpt bihormonal cells, 3 for sst1.1GFP^high^, 3 for sst1.1GFP^low^). Reads were mapped and aligned to the zebrafish genome GRCz11 from Ensembl gene annotation version 92 using STAR(Dobin et al., 2013). Gene expression levels were calculated with featureCounts (http://bioinf.wehi.edu.au/featureCounts/) and differential expression determined with DESeq2(Love, Huber, & Anders, 2014). Expression values are given as normalized read counts. Poorly expressed genes with mean normalized expression counts <10 were excluded from the subsequent analyses. DESeq2 uses Wald test for significance with posterior adjustment of P values (Padj) using Benjamini and Hochberg multiple testing. The differentially expressed (DE) genes identified with a Padj cutoff of 0.05 and fold change above 2 were submitted for GO analysis using WebGestalt tool (Liao, Wang, Jaehnig, Shi, & Zhang, 2019).

The genes enriched in β-cells and *sst2* δ-cells above 4-fold were identified using sequences obtained previously (Tarifeño-Saldivia et al., 2017) with prior mapping on the more recent GRCz11 v92 assembly of the zebrafish genome; they thus slightly differ from the gene list previously published (provided in Figure3-Source Data 2). Then, new enrichment was recalculated to take into account the new transcriptomic data obtained for *sst1.1* δ-cells from *Tg(sst1.1:eGFP)* and the new β-cells from *Tg(ins:NTR*-P2A-mCherry*) (presented in Figure 4-Source Data 3).

### Statistical analyses

Graphs and statistical analyses were performed using GraphPad Prism 8 and means ± SD are shown; the statistical tests are described in the legend of the Figures.

## Acknowledgments

The authors thank the GIGA technology platforms GIGA-Zebrafish, GIGA-Genomics and GIGA-Imaging. The authors also thanks Chris Wright for providing the Pdx1 antibody.

## Figure Supplements legends

**Figure 1-figure supplement 1. *Tg(sst1.1:GFP)* is active in sst1.1+ cells and not in β-cells**

A) *In situ* hybridization on 3 dpf *Tg(sst1.1:GFP)* embryo using a *sst1.1* antisense RNA probe (red) combined with immunodetection of the GFP protein (green) revealing co-localization between endogenous *sst1.1* transcripts and GFP cells.

B) Immunofluorescence in adult *Tg(sst1.1:GFP); Tg(ins:NTR*-P2A-mCherry)* islet showing co-localization between GFP (green) and the endogenous SST protein (red) and not with mCherry β-cells (grey).

**Figure 1-figure supplement 2. Analysis of sst1.1:GFP and *ins:NTR*-P2A-mCherry* fluorescent cells by flow cytometry**

GFP and mCherry fluorescence analysis by flow cytometry of dissociated main islets (3-4 pooled islets) isolated from *Tg(sst1.1:GFP);Tg(ins:NTR*-P2A-mCherry)^ulg034^* adult fish. Representative plots showing fluorescent cells along GFP and mCherry axes. The populations of interest are delimited with dashed lines that highlight the gates that were used for cell sorting. The positive/negative fluorescence thresholds delineate quadrants that were considered for fluorescent cells quantification.

**Figure 2-figure supplement 1. Bihormonal cell formation following β-cell ablation with Diphteria Toxin A**

β-cell ablation performed using the cytotoxic Diphteria Toxin chain A (DTA) inducible system in *Tg(ins:loxP-mCherry-loxP-DTA); Tg(ins:CRE-ERT2)*. 7 dpf larvae were treated with 4-OHT to trigger the recombination of the loxP-mCherry-loxP cassette and allow DTA expression and β-cells death, and then analysed 9 days after by immunofluorescence. Like in the NTR/prodrug system, DTA induces the formation of INS+ SST+ bihormonal cells.

**Figure 2-figure supplement 2. Experimental design for the β-cell lineage tracing after ablation**

β-cell lineage tracing was performed in Tg(ins:CRE-ERT2); Tg(ubb:loxP-CFP-loxP-zsYellow); Tg(sst1.1:GFP); Tg(ins:NTR-P2A-mCherry) larvae as presented in Figure 2C-F.

**Figure 3-figure supplement 1. Gcg is not detected in bihormonal cells**

Immunodetection of GFP (green), mCherry (red) and GCG (grey) adult *Tg(sst1.1:GFP); Tg(ins:NTR*-P2A-mCherry)* principal islets in CTL and 20 dpt conditions showing bihormonal (GFP+ mCherry+) cells at 20 dpt and non-overlapping GCG staining (white arrowheads).

**Figure 4-figure supplement 1. sst1.1:GFP cells delineate two distinct δ-cell subpopulations**

A) Fluorescence analysis by flow cytometry of GFP+ mCherry-cells from *Tg(sst1.1:eGFP); Tg(ins:NTR*-P2A-mCherry)* islets. Two populations, namely GFP^high^ and GFP^low^, can be identified based on their GFP intensity.

B) Heatmap representation of the transcriptomes of GFP^high^ and GFP^low^ cells (3 replicates each) (significant DE genes).

C) In situ hybridization on 3 dpf *Tg(sst1.1:GFP)* embryo using a *sst2* antisense RNA probe (red) combined with immunodetection of the GFP protein (green) revealing co-localization between endogenous *sst2* transcripts and GFP^low^ + cells. In contrast, GFP^high^ cells do not present detectable transcripts of *sst2*.

## List of Figure-Source Data

Figure 3-Source Data 1

Figure 3-Source Data 2

Figure 3-Source Data 3

Figure 4-Source Data 1

Figure 4-Source Data 2

Figure 4-Source Data 3

Figure 4-Source Data 4

Figure 4-Source Data 5

Figure 5-Source Data 1

Figure 5-Source Data 2

Figure 5-Source Data 3

