## Supplementary figures and images for "A δ-cell subpopulation with a pro-β-cell identity confers efficient age-independent recovery in a zebrafish model of diabetes"

### Figure 1 Supplement figure 1

Figure 1-figure supplement 1

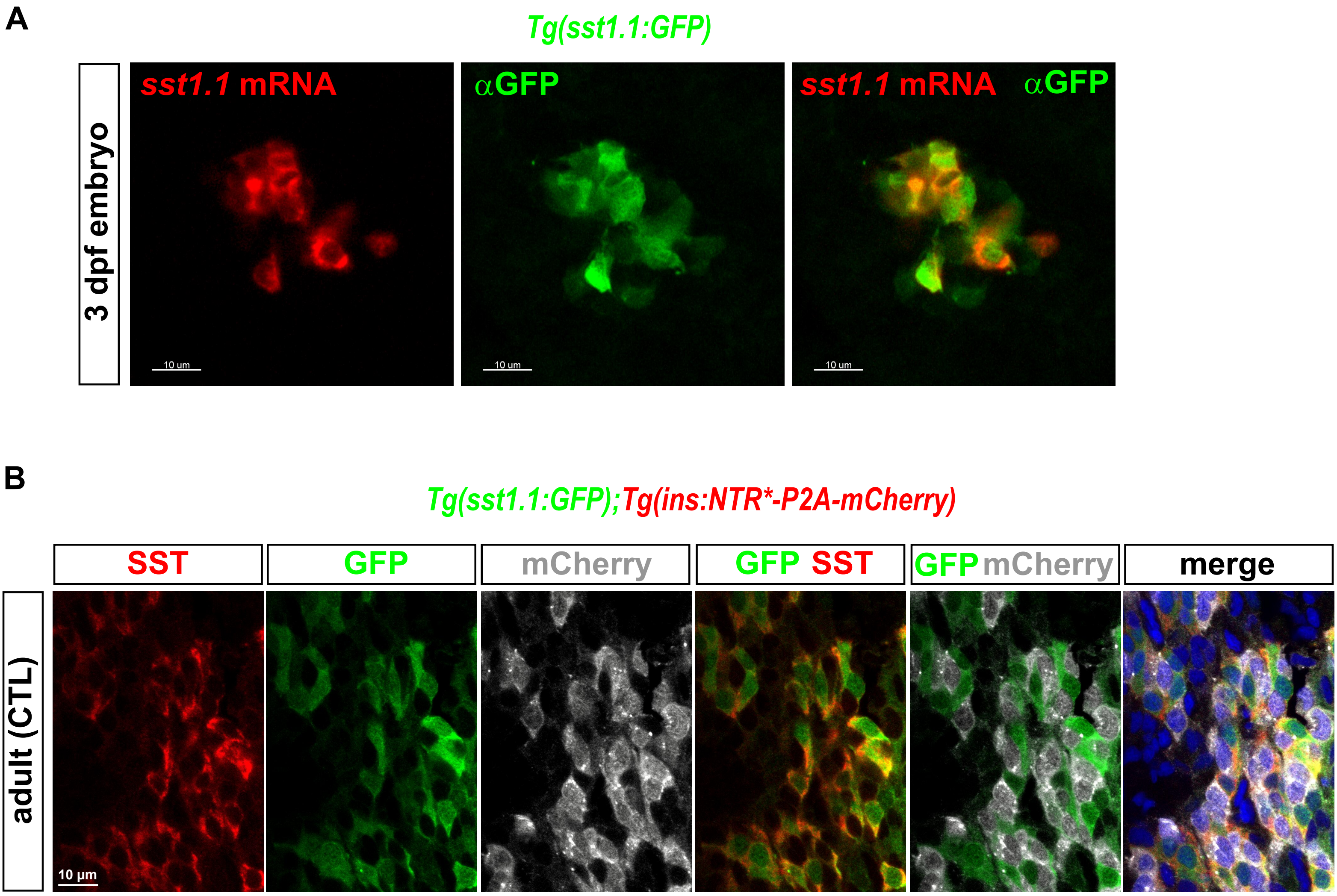

### Figure 1 Supplement figure 2

#### Figure 1-figure supplement 2

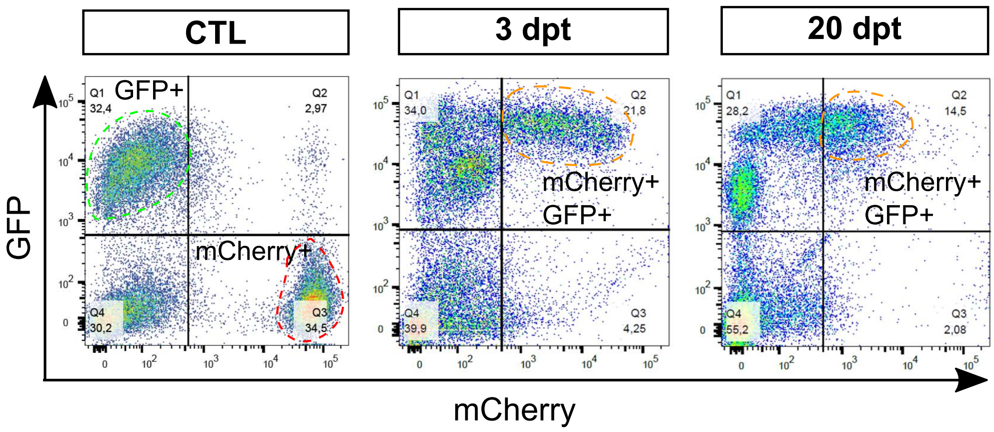

### Figure 2 Supplement figure 1

#### Figure 2-figure supplement 1

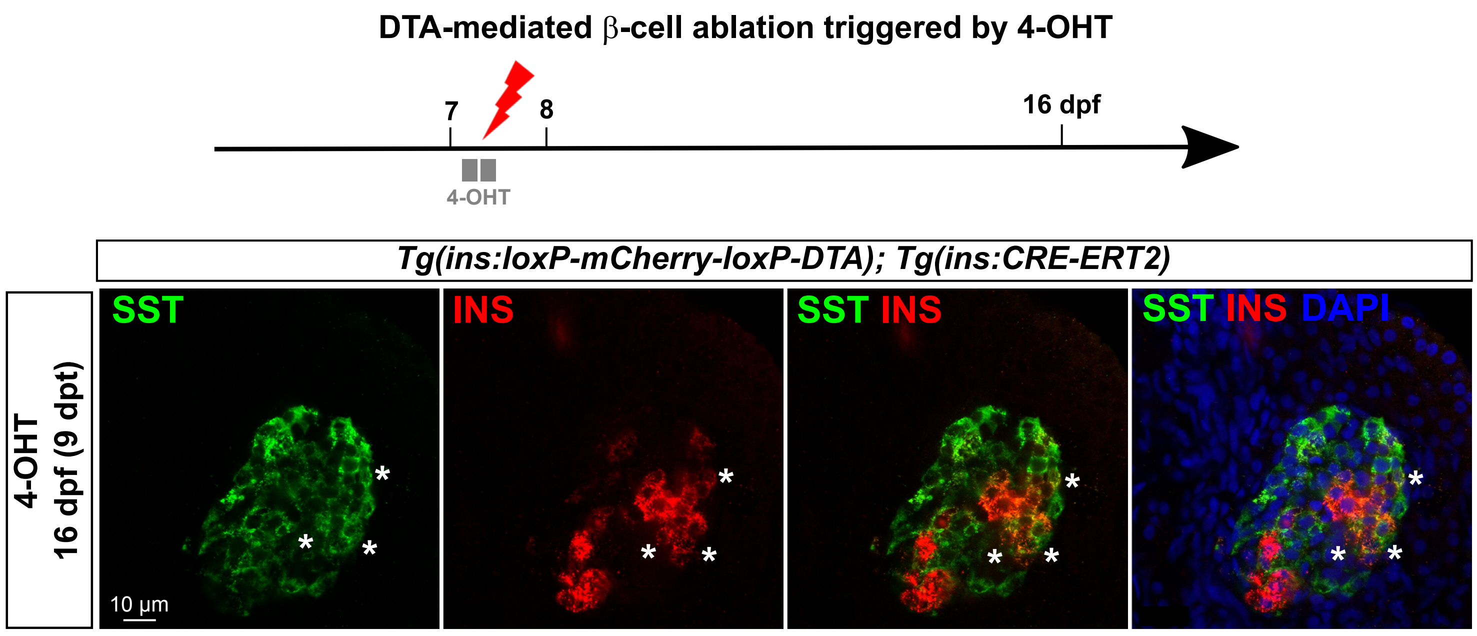

### Figure 2 Supplement figure 2

**Figure 2-figure supplement 2**

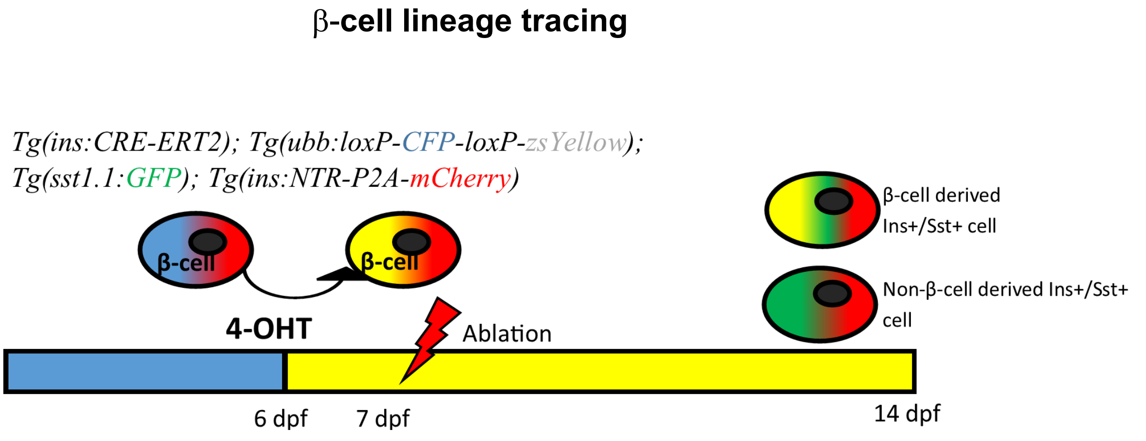

### Figure 3 Supplement figure 1

**Figure 3-figure supplement 1**

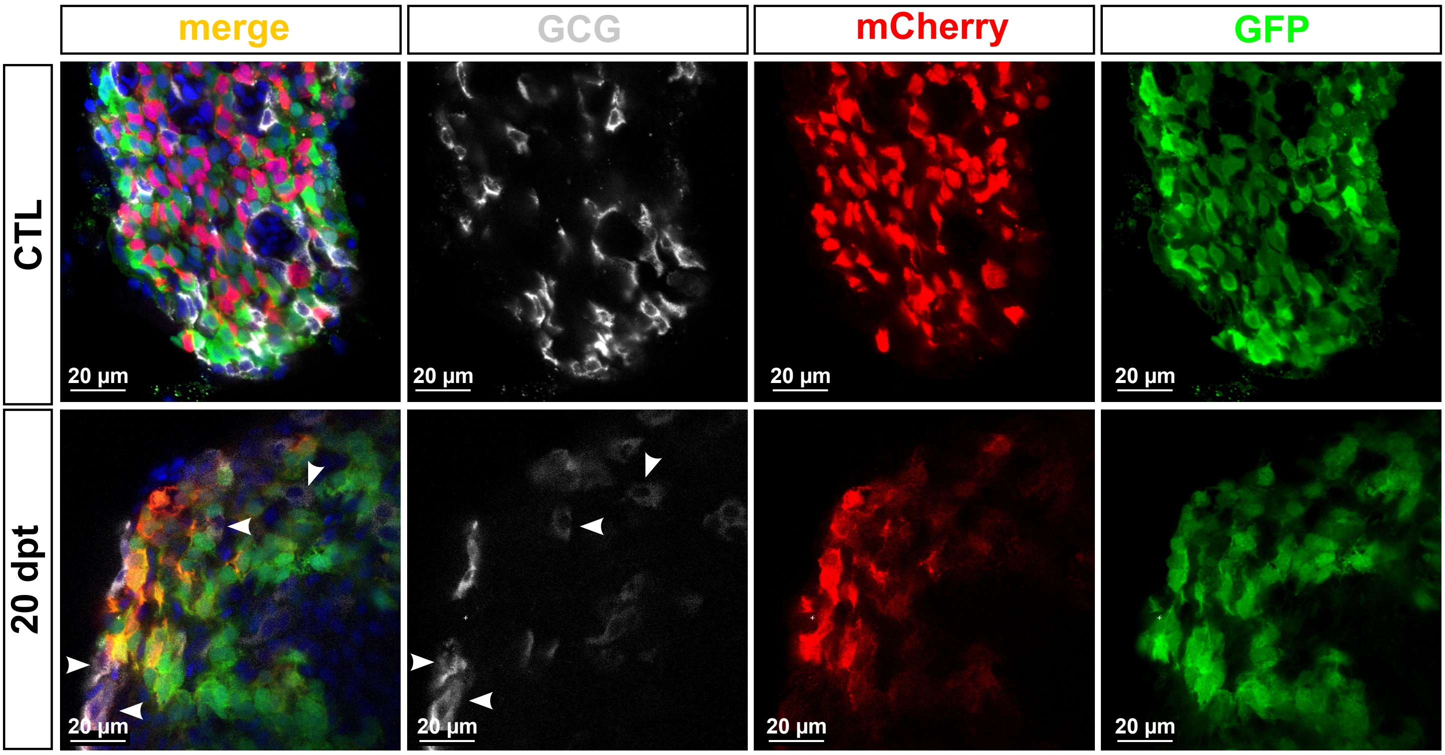

### Figure 4 Supplement figure 1

#### Figure 4-figure supplement 1

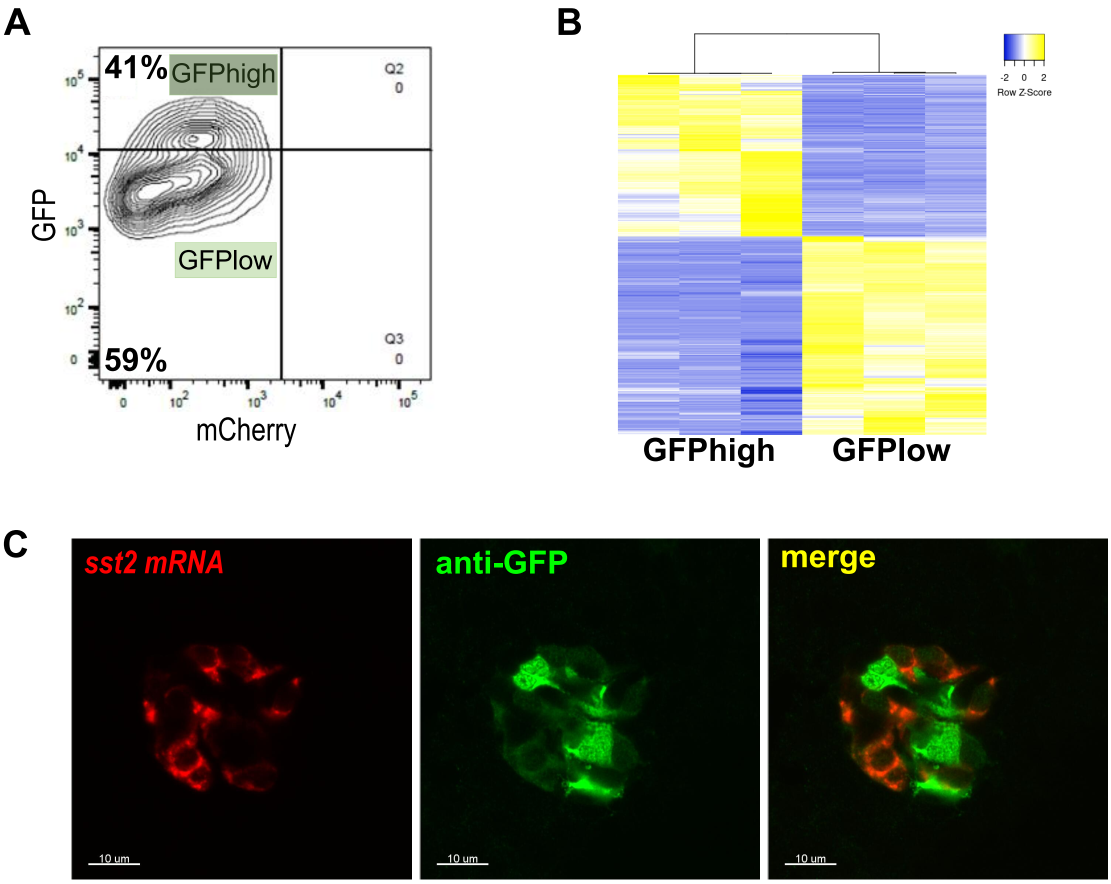
